# A genome-wide screen in macrophages identifies PTEN as required for myeloid restriction of *Listeria monocytogenes* infection

**DOI:** 10.1101/2022.12.12.520030

**Authors:** Rochelle C. Glover, Nicole H. Schwardt, Shania-Kate E. Leano, Madison E. Sanchez, Maureen K. Thomason, Andrew J. Olive, Michelle L. Reniere

## Abstract

*Listeria monocytogenes* (*Lm*) is an intracellular foodborne pathogen which causes the severe disease listeriosis in immunocompromised individuals. Macrophages play a dual role during *Lm* infection by both promoting dissemination of *Lm* from the gastrointestinal tract and limiting bacterial growth upon immune activation. Despite the relevance of macrophages to *Lm* infection, the mechanisms underlying phagocytosis of *Lm* by macrophages are not well understood. To identify host factors important for *Lm* infection of macrophages, we performed an unbiased CRISPR/Cas9 screen which revealed pathways that are specific to phagocytosis of *Lm* and those that are required for internalization of bacteria generally. Specifically, we discovered the tumor suppressor PTEN promotes macrophage phagocytosis of *Lm* and *L. ivanovii*, but not other Gram-positive bacteria. Additionally, we found that PTEN enhances phagocytosis of *Lm* via its lipid phosphatase activity by promoting adherence to macrophages. Using conditional knockout mice lacking *Pten* in myeloid cells, we show that PTEN-dependent phagocytosis is important for host protection during oral *Lm* infection. Overall, this study provides a comprehensive identification of macrophage factors involved in regulating *Lm* uptake and characterizes the function of one factor, PTEN, during *Lm* infection *in vitro* and *in vivo*. Importantly, these results demonstrate a role for opsonin-independent phagocytosis in *Lm* pathogenesis and suggest that macrophages play a primarily protective role during foodborne listeriosis.

**Author Summary:** *Listeria monocytogenes* (*Lm*) is a bacterial pathogen that causes the foodborne illness listeriosis primarily in immunocompromised, elderly, and pregnant individuals. Listeriosis is one of the deadliest bacterial infections known, with a mortality rate of ~30% even when treated with antibiotics. The high mortality rate of listeriosis is due to inefficient restriction of *Lm* by the immune system, and subsequent spread of bacteria beyond the gastrointestinal tract to internal organs such as the liver and brain. Macrophages are important for immune clearance of *Lm* but are also hypothesized to promote dissemination of intracellular *Lm*; thus, studies of *Lm*-macrophage interactions are critical for understanding the balance between bacterial growth and restriction by these phagocytes. We performed a forward genetic screen in macrophages and discovered that the tumor suppressor PTEN promotes phagocytosis of *Lm* by enhancing adherence to macrophages. These results demonstrate a novel function of macrophage PTEN, which canonically acts as a repressor of phagocytosis. In addition, we found that PTEN protects mice from severe disease and lowers bacterial burdens following oral inoculation of *Lm*. Our results demonstrate for the first time that macrophage phagocytosis is an important immune defense against invasive *Lm* during the foodborne route of infection.

## Introduction

The genus *Listeria* consists of 21 species of Gram-positive bacteria, of which only two, *Listeria monocytogenes* (*Lm*) and *L. ivanovii*, are considered pathogenic to mammals (1). *Lm* is a major human pathogen and etiologic agent of the severe foodborne disease listeriosis, which progresses to meningitis in immunocompromised and elderly patients, or neonatal septicemia and abortion in pregnant individuals (2). Following consumption of *Lm*-contaminated food, the bacteria colonize the gastrointestinal (GI) tract and cross the intestinal epithelium into the underlying lamina propria where they encounter resident immune cells and replicate intracellularly (3). From there, *Lm* disseminates through the lymphatics to the mesenteric lymph nodes (MLN) and spleen, and through the bloodstream to the liver and gallbladder (4). After replicating to high numbers in these organs, the bacteria re-enter systemic circulation and cross the blood-brain barrier or placental barrier to initiate central nervous system (CNS) infection or fetal infection, respectively (5). While the route of *Lm* dissemination has been characterized, the precise mechanisms underlying the method of dissemination from the GI tract are a current area of study.

A growing body of literature supports a model in which phagocytic monocytes and macrophages promote dissemination of *Lm* to internal organs. Development of an oral infection model of listeriosis in recent years has allowed for closer examination of *Lm* dissemination from the lamina propria, with emerging hypotheses suggesting transit of bacteria from the lamina propria to the MLN occurs within phagocytes (3,4). In support of this model, intracellular replication in growth-permissive lamina propria cells, such as resident macrophages, is important for initial dissemination to the MLN (6). In addition, several studies have demonstrated the importance of *Lm*-infected monocytes in establishing CNS infection in both intravenous (i.v.) and oral models of infection (7–11). In contrast to the role macrophages and monocytes play in systemic dissemination of *Lm*, these inflammatory phagocytes are also critical for bacterial clearance and host protection (12,13). Growth-permissive macrophages become listericidal upon activation by interferon gamma (IFNγ), which is crucial for limiting bacterial replication *in vivo* (14–16). Importantly, studies demonstrating the role of macrophages in host protection against *Lm* were performed using i.v. inoculation, which models systemic infection but bypasses the intestinal phase. Thus, the relative contributions of macrophages to pathogen dissemination versus bacterial restriction during foodborne listeriosis remain unclear, particularly in the case of lamina propria macrophages during the GI phase of infection.

As an intracellular pathogen, *Lm* invades both phagocytic and non-phagocytic cells and undergoes an intracellular lifecycle involving escape from the endosomal vacuole, replication in the host cell cytosol, and intercellular spread using actin-based motility (17). Research over the last 30 years has revealed the mechanisms of *Lm* invasion of non-phagocytic cells, including the host receptors, bacterial invasins, and downstream signaling components involved (5). In contrast, relatively little is known about the host and bacterial factors involved in uptake of *Lm* by professional phagocytes, despite their importance to *Lm* pathogenesis. Existing studies on *Lm* entry into phagocytes have primarily focused on complement-mediated phagocytosis, which restricts *Lm* to the bactericidal phagocytic vacuole (18,19). However, replication of *Lm* in the cytosol of naïve macrophages occurs *in vitro* and *in vivo*, indicating the presence of biologically relevant opsonin-independent phagocytosis pathways. Early *in vitro* work demonstrated that phagocytosis of non-opsonized *Lm* uses distinct receptors and results in cytosolic infection, however this pathway was not characterized or subsequently investigated (20,21).

In this study, we sought to identify macrophage factors involved in mediating cytosolic infection by *Lm*. Using a flow cytometry-based approach, we performed a genome-wide CRISPR/Cas9 knockout screen in macrophages for mutant cells resistant to *Lm* infection. Our screen identified biological pathways involved in phagocytosis of *Lm* specifically, as well as Gram-positive bacteria broadly. We found that the phosphatase and tumor suppressor PTEN (Phosphatase and Tensin homolog) was specifically required for uptake of *Lm* by human and mouse macrophages. PTEN promotes phagocytosis of *Lm* by enhancing adherence to macrophages and uptake through this pathway is more efficient than PTEN-independent mechanisms. Tight control of phosphatidylinositol (3,4,5)-triphosphate (PIP_3_) signaling regulates phagocytosis of *Lm*, as both PIP_3_ production and turnover by PI3K and PTEN, respectively, are required for uptake. Using an oral murine listeriosis model in conditional *Pten* knockout mice, we demonstrate that myeloid PTEN is dispensable for dissemination from the GI tract and restricts *Lm* replication *in vivo*. Our unbiased forward genetic screen in macrophages constitutes a major step towards comprehensively identifying the host determinants of *Lm* infection and reveals a novel function of PTEN in promoting phagocytosis of a bacterial pathogen.

## Results

### CRISPR/Cas9 screen reveals host cell determinants of macrophage infection

To begin investigating phagocytosis of non-opsonized *Lm*, we first assessed uptake of various *Listeria* species using gentamicin protection assays. Primary murine bone marrow-derived macrophages (BMDMs) were infected, followed immediately by short centrifugation to control for any motility differences between species. Gentamicin was added 30 minutes post-infection to kill extracellular bacteria and intracellular bacteria were enumerated 1 hour post-infection. *Lm* was internalized more efficiently than either *L. ivanovii* or *L. innocua*, with *L. innocua* being taken up only ~15% as efficiently as *Lm* (Figure 1A). These results suggest distinct mechanisms of phagocytosis between *Listeria* species under non-opsonizing conditions, although the bacterial and host factors involved in this differential uptake are unknown.

**Fig 1.**
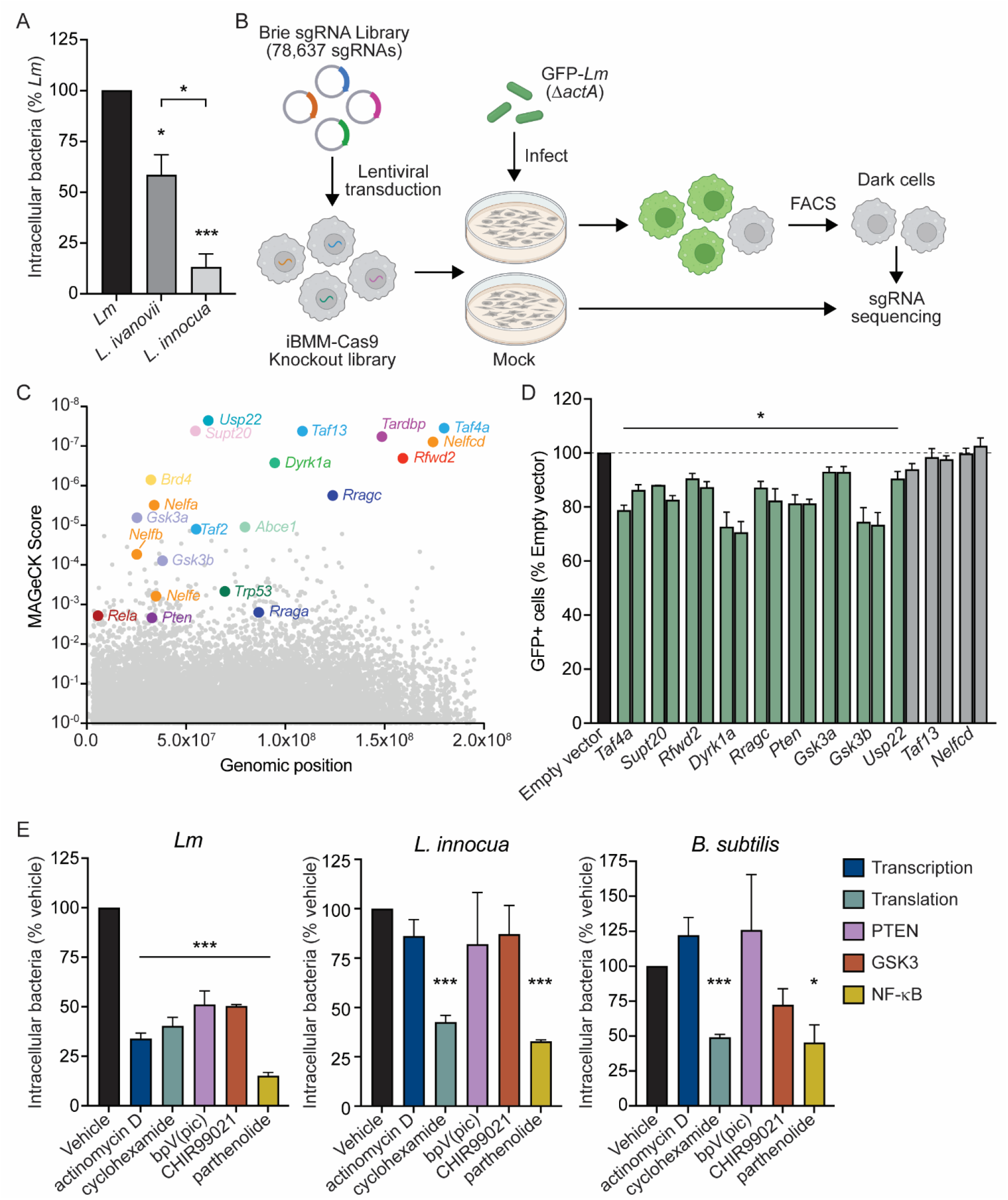
Genome-wide CRISPR/Cas9 screen identifies novel determinants of *Lm* macrophage infection. (A) Gentamicin protection assay measuring bacterial uptake by BMDMs 1 hour post-infection. Bacteria (MOI=1) were centrifuged onto BMDMs at 300 x *g* for 3 minutes prior to incubation. Data are normalized to uptake of *Lm*. (B) Schematic of the CRISPR/Cas9 screen in iBMMs. Three independent iBMM knockout libraries were generated using the genome-wide Brie sgRNA library. Cells were infected with mock or GFP-*Lm* at MOI=10 for 4 hours. 6 hours post-infection, uninfected cells (GFP^-^) were isolated by FACS. sgRNA abundances in sorted cells versus mock-infected controls were measured by Illumina sequencing and MAGeCK analysis. (C) MAGeCK score of each gene in the Brie library from three independent screen replicates. Gene names are depicted for the top 14 hits (lowest MAGeCK scores), several related genes, and specific genes identified by Metascape analysis. (D) Validation of CRISPR/Cas9 screen. iBMMs were transduced with empty vector or sgRNAs targeting selected hits. For each gene, two independent sgRNAs were used to generate two knockout cell lines. Cell lines were infected with GFP-*Lm* as in (B) and analyzed by flow cytometry. Data are normalized to iBMMs transduced with empty vector. Non-significant results are depicted in gray. (E) Gentamicin protection assay measuring bacterial uptake by iBMMs in the presence of 10 μg/mL actinomycin D, 1 μg/mL cycloheximide, 5 μM bpV(pic), 10 μM CHIR99021, or 10 μM parthenolide. All data displayed are the means and standard error of the mean (SEM) of at least three biological replicates. \**p*<0.05, \*\*\**p*<0.001 as determined by unpaired *t* tests. *p* values for significant results in (D) ranged from <0.0001 to 0.025.

To identify macrophage factors required for phagocytosis of *Lm*, we performed a genome-wide CRISPR/Cas9 knockout screen in immortalized bone marrow-derived macrophages expressing Cas9 (iBMM) (22). Similar to BMDMs, iBMMs exhibited enhanced uptake of *Lm* as compared to other *Listeria* species (S1A Fig). Three independent iBMM knockout libraries were generated using the murine Brie sgRNA library (23). To select for iBMM mutants defective for *Lm* uptake, we constructed a reporter strain of *Lm* constitutively expressing GFP (GFP-*Lm*) in an *actA-*deficient background to eliminate intercellular spread between different mutant cells. Using standard infection conditions of a multiplicity of infection (MOI) of 1 for 1 hour resulted in only ~5% of macrophages becoming GFP^+^ (S1B Fig). To overcome this bottleneck and maintain coverage of the knockout library, each iBMM knockout library was infected with GFP-*Lm* using an MOI of 10 for 4 hours, which resulted in approximately 80% GFP^+^ iBMMs (S1B Fig). 6 hours post-infection, phagocytosis-deficient (*i.e*. GFP^−^) cells were isolated using fluorescence-activated cell sorting (FACS). sgRNAs were amplified from sorted cells and mock-infected control libraries and abundance in each population was measured by Illumina sequencing (Figure 1B). Importantly, 1000-fold coverage of the library was maintained at each step, including transduction, passaging, and sorting.

Using Model-based Analysis of Genome-wide CRISPR/Cas9 Knockout (MAGeCK) (24), we identified 557 sgRNAs enriched in the GFP^-^ populations compared to the mock-infected libraries (*p* < 0.01), corresponding to 235 genes (Figure 1C, S1 Table). Initial examination of these candidates found several sets of genes belonging to the same protein family or protein complex: TFIID complex proteins (*Taf1d, Taf2, Taf3, Taf4a, Taf5, Taf8, Taf10*, and *Taf13*); SAGA complex proteins (*Usp22, Supt7l, Supt20, Taf5l, Taf6l, Atxn7l3, Tada2b*, and *Tada1*); negative elongation factor (*Nelfa, Nelfb, Nelfcd*, and *Nelfe*); Ras-related GTP-binding proteins (*Rraga* and *Rragc*); glycogen synthase kinase 3 (*Gsk3a* and *Gsk3b*); and translation initiation factors (*Eif3h, Eif4g2*, and *Eif5*). Identification of multiple independent genes within the same complex suggests that our genome-wide screen was robust. In addition, we performed gene ontology analysis using Metascape to identify biological pathways enriched in our data set (25). Several enriched pathways were involved in chromatin remodeling, mRNA transcription, and mRNA processing (S1C Fig), suggesting that proper host cell transcription may be required for phagocytosis of *Lm*. Interestingly, we also identified signaling pathways involving major regulators of cell proliferation and survival, including PTEN (*Pten*), p53 (*Trp53*), and NF-κB (*Rela*) (Figure 1C and S1C Fig). Overall, analysis of the screen provides confidence in our approach and indicates that we have identified several pathways that have not been previously implicated in *Lm* uptake.

To validate the screen, individual knockout cell lines of a subset of genes were generated using 2 sgRNAs per gene and genome editing was confirmed using Inference of CRISPR Edits (ICE) analysis (26). Knockout iBMMs were infected with GFP-*Lm* in the same manner as the original screen, and GFP^+^ cells were quantified using flow cytometry. Of the eleven candidate genes tested, eight exhibited significantly reduced intracellular *Lm* with both sgRNAs and one with a single sgRNA (Figure 1D). The two candidate genes which had no change in *Lm* uptake with either sgRNA (*Nelfcd* and *Taf13*) had <30% CRISPR-InDel frequency, as measured by ICE analysis, or were unable to be confirmed by sequencing. Together, these data validate the power of our screen to identify novel genes involved in *Lm* macrophage infection.

Based on the length of infection used in the screen, we anticipated the possibility of identifying genes involved not only in phagocytosis but also in *Lm* vacuolar escape and cytosolic growth. In addition, we expected to identify genes required for the general process of phagocytosis along with those specific for internalization of *Lm*. To test uptake more directly and assess the specificity of the identified pathways, gentamicin protection assays were used to measure internalization of *Lm*, *L. innocua*, and the related Gram-positive bacterium *Bacillus subtilis*. As an orthologous approach to genetic knockouts, we used chemical inhibitors targeting transcription (actinomycin D), translation (cycloheximide), PTEN [bpV(pic)], GSK3 (CHIR99021), and NF-κB (parthenolide). All inhibitors tested blocked internalization of *Lm* during the 1-hour infection, suggesting the involvement of these pathways in uptake of *Lm* rather than later stages of infection (Figure 1E). Interestingly, inhibitors targeting transcription, PTEN, and GSK3 had no effect on uptake of *L. innocua* or *B. subtilis*, indicating their specificity for internalization of *Lm*. We also found that host cell translation and NF-κB were required for uptake of all three species (Figure 1E). Collectively, these data demonstrate the breadth of our screen in identifying regulators of phagocytosis both specific to *Lm* and involved more broadly in phagocytosis of bacteria.

### PTEN is required for uptake of *Lm* by macrophages

Given that *Lm* is internalized by macrophages using distinct mechanisms (Figure 1A), we aimed to further characterize pathways which uniquely regulate *Lm* uptake. Metascape analysis identified the PTEN regulation pathway as enriched in our dataset (S1C Fig and Figure 2A), and two members of this pathway, GSK3 and PTEN itself, were both required specifically for phagocytosis of *Lm* by macrophages (Figure 1E). Therefore, we investigated the involvement of PTEN in *Lm* uptake. PTEN (<u>P</u>hosphatase and <u>Ten</u>sin homolog) is a dual-specificity phosphatase primarily known for its tumor suppressor activity (27). PTEN blunts the PI3K signaling cascade, halting the promotion of cell survival and proliferation by this pathway (28,29). While the role of PTEN in cancer biology has been very well-studied, relatively little is known about the role PTEN might play in the regulation of bacterial phagocytosis by macrophages.

**Fig 2.**
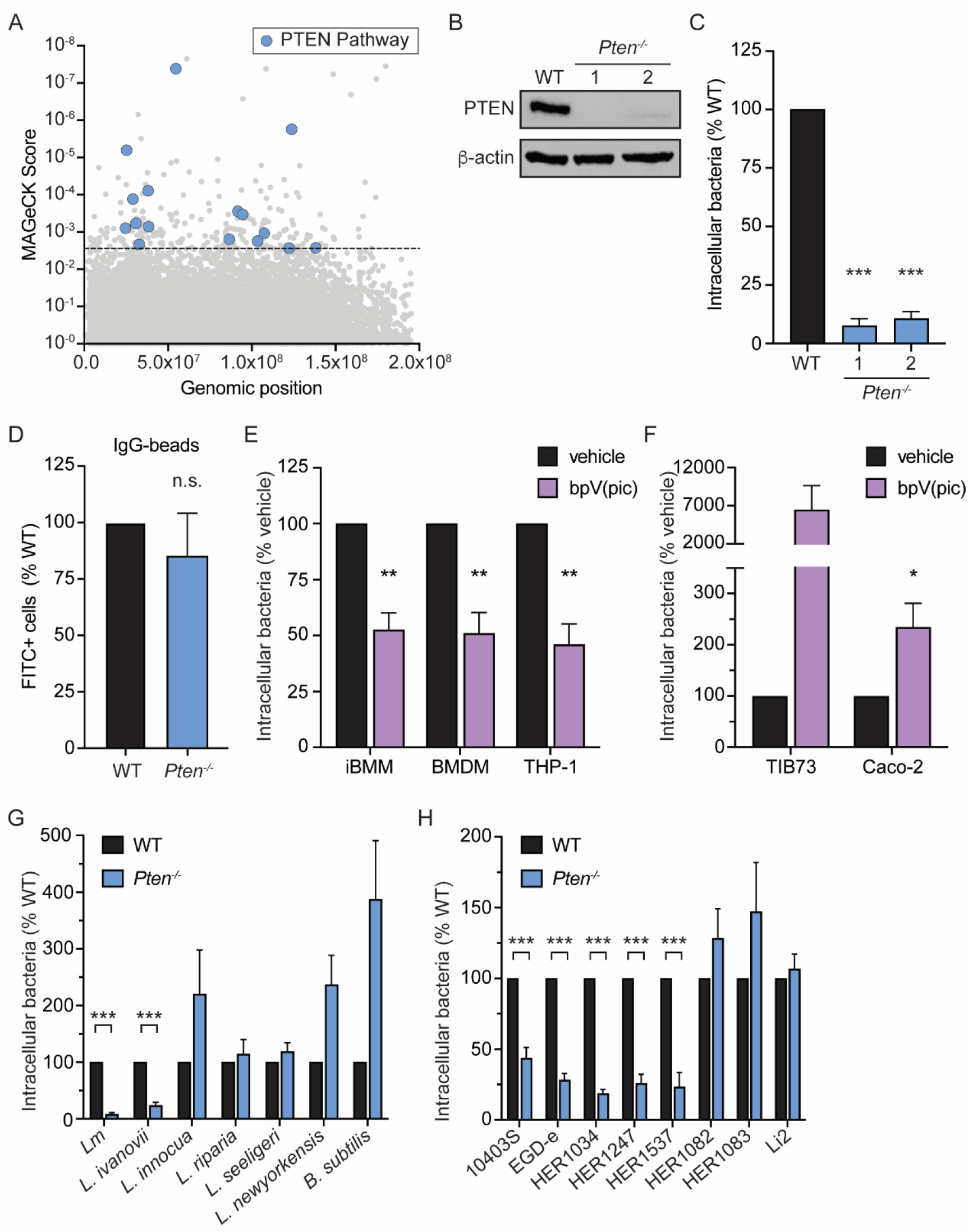
PTEN is a novel host factor which enhances phagocytosis of specific *Lm* strains. (A) Hits from CRISPR/Cas9 screen belonging to the PTEN regulation pathway (n=16) as identified by Metascape analysis. Dashed line indicates *p*-value cutoff (*p*<0.01). (B) Immunoblot of PTEN in WT and *Pten*^−/−^ iBMM clones. β-actin is used as a loading control. (C) Gentamicin protection assay measuring uptake of *Lm* by *Pten*^−/−^ iBMMs. Two independent single-cell clones were analyzed. Data are normalized to WT iBMMs. (D) Uptake of IgG-opsonized FITC-labeled latex beads by WT and *Pten*^−/−^ iBMMs. Beads were added to cells for 30 minutes and uptake was quantified by flow cytometry. (E) Gentamicin protection assay measuring uptake of *Lm* by iBMMs, BMDMs, and PMA-differentiated THP-1 cells. Cells were treated with vehicle or 5 μM bpV(pic). Data are normalized to vehicle-treated cells. (F) Gentamicin protection assay measuring uptake of *Lm* by TIB73 and Caco-2 cells. Cells were treated with vehicle or 5 μM bpV(pic). Data are normalized to vehicle-treated cells. (G) Gentamicin protection assay measuring uptake of bacteria by WT and *Pten*^−/−^ iBMMs. Data are normalized to WT iBMMs for each species. (H) Gentamicin protection assay measuring uptake of *Lm* strains by WT and *Pten*^−/−^iBMMs. Data are normalized to WT iBMMs for each strain. All data are means and SEM of at least three biological replicates. \**p*<0.05, \*\**p*<0.01, \*\*\**p*<0.001, n.s. = non-significant, as determined by unpaired *t* tests.

To elucidate the role of PTEN in the context of *Lm* infection, we first derived single-cell clones from the iBMM *Pten* CRISPR knockout population (Figure 1D) and confirmed the absence of PTEN protein by immunoblot for two independent clones (Figure 2B). A small amount of PTEN protein at a lower molecular weight was detected for Clone 2, however sequencing analysis revealed a 45 base pair deletion in the phosphatase domain of this clone. Each cell line was infected with *Lm* and uptake was assessed by gentamicin protection assay. Compared to wildtype (WT) iBMMs, *Pten*^−/−^ iBMMs exhibited a dramatic reduction in *Lm* uptake (Figure 2C), consistent with chemical inhibition of PTEN. As both clones had similar phenotypes, iBMM *Pten*^−/−^ Clone 1 was used for the remainder of our experiments. To assess the overall phagocytic ability of *Pten*^−/−^ iBMMs, we measured phagocytosis of fluorescently-labeled latex beads, an established method for studying basal phagocytic activity(30). IgG-opsonized FITC-labeled beads were added to WT and *Pten*^−/−^ iBMMs for 30 minutes, after which excess beads were removed. Fluorescence of any remaining extracellular beads was quenched with trypan blue and FITC fluorescence was quantified by flow cytometry. Phagocytosis of fluorescent beads was not significantly different between WT and *Pten*^−/−^ iBMMs (Figure 2D), indicating that basal phagocytic activity was not altered by loss of *Pten*. In addition, these results suggest that PTEN is not required for Fcγ receptor-mediated phagocytosis in iBMMs.

We next investigated the role of PTEN in *Lm* internalization in various cell types. iBMMs, BMDMs, and THP-1 human monocyte-derived macrophages were treated with the well-characterized potent competitive PTEN inhibitor bpV(pic) (31). Uptake of *Lm* was reduced to similar levels in all three macrophage cell types tested (Figure 2E), indicating that regulation of *Lm* uptake by PTEN is conserved in primary murine macrophages as well as human macrophages. *Lm* also induces phagocytosis-like internalization into non-phagocytic cells (32). In contrast to macrophages, TIB73 murine hepatocytes and Caco-2 human colonic epithelial cells exhibited increased *Lm* internalization following treatment with bpV(pic) (Figure 2F). These data demonstrate that PTEN is required for phagocytosis of *Lm* specifically by macrophages.

Chemical inhibition of PTEN specifically blocked uptake of *Lm* but not *L. innocua* or *B. subtilis* (Figure 1E). To confirm the specificity of PTEN for regulation of *Lm* uptake, we assessed phagocytosis of various *Listeria* species and *B. subtilis* in WT and *Pten*^−/−^ iBMMs. Consistent with our previous results, *Pten*^−/−^ iBMMs were deficient for uptake of the pathogenic species *Lm* and *L. ivanovii*, but not the non-pathogens *L. innocua*, *L. riparia*, *L. seeligeri*, *L. newyorkensis*, or *B. subtilis* (Figure 2G). Given that *Lm* and *L. ivanovii* are taken up more efficiently by macrophages than *L. innocua*, we compared uptake efficiency in WT and *Pten*^−/−^ iBMMs between these species by determining the percentage of the initial inoculum that was taken up under each condition. Uptake efficiency of *Lm* and *L. ivanovii* by *Pten*^−/−^ iBMMs was reduced to that of *L. innocua* in WT cells (S2 Fig). These data suggest that the enhanced macrophage uptake of *Lm* and *L. ivanovii* relative to *L. innocua* observed in Figure 1A is PTEN-dependent. In addition to assessing uptake of different species, we also measured phagocytosis of various *Lm* strains in *Pten*^−/−^ iBMMs. Similar to the lab strain 10403S used in all prior experiments, four other *Lm* strains exhibited reduced uptake in *Pten*^−/−^ iBMMs (Figure 2H). In contrast, uptake of three *Lm* strains (HER1082, HER1083, and Li2) remained unchanged in *Pten*^−/−^ iBMMs. Similar to our species-level analysis, *Lm* strains which required PTEN for uptake were internalized more efficiently by macrophages, and this advantage was lost in *Pten*^−/−^ iBMMs (S2 Fig). Overall, these data demonstrate that PTEN enhances uptake of specific species and *Lm* strains by macrophages.

### PTEN promotes adherence of *Lm* to macrophages

Given that *Pten*^−/−^ iBMMs have reduced intracellular CFU as early as 1 hour post-infection, we hypothesized that PTEN is not involved in escape from the vacuole. To directly test this, we compared uptake of wildtype *Lm* and a listeriolysin O-deficient strain (Δ*hly*), which is unable to escape the phagocytic vacuole. If the defect in *Pten*^−/−^ iBMMs was due to impaired vacuolar escape, we would expect to see no difference in intracellular CFU between WT and *Pten*^−/−^ iBMMs during infection with the Δ*hly* mutant that is trapped in the vacuole. However, the Δ*hly* strain displayed a similar reduction in CFU in *Pten*^−/−^ iBMMs compared to wildtype *Lm* (Figure 3A), indicating that PTEN does not play a role in vacuolar escape of *Lm*. In addition, there was no difference in the intracellular growth rate of wildtype *Lm* between WT and *Pten*^−/−^ iBMMs (Figure 3B). Taken together, these data confirm that while PTEN promotes internalization of *Lm*, PTEN is not important for *Lm* escape from the vacuole or replication in the host cytosol.

**Fig 3.**
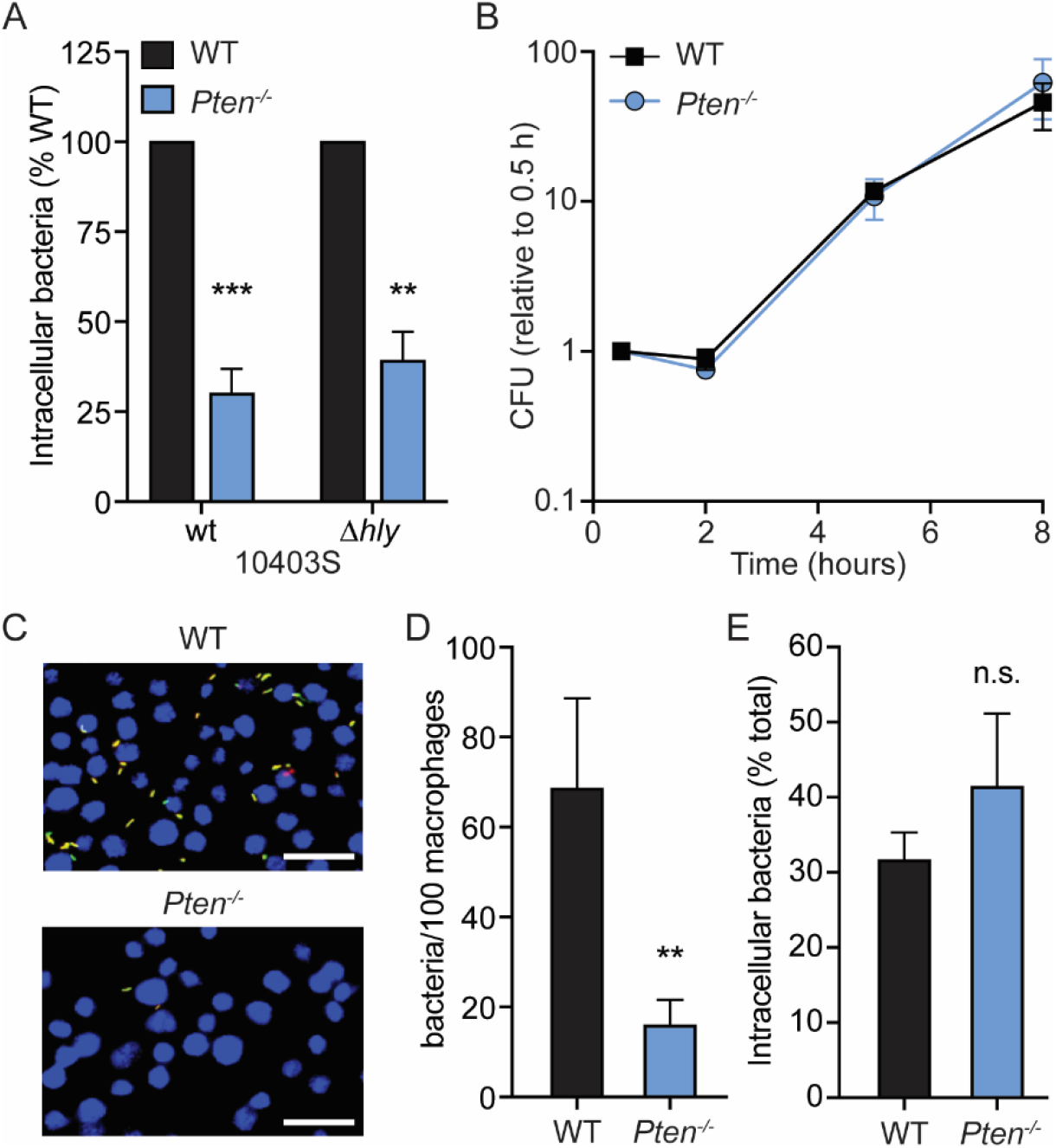
PTEN promotes adherence of *Lm* to macrophages. (A). Gentamicin protection assay measuring uptake of *Lm* wildtype (wt) and Δ*hly* strains by WT and *Pten*^−/−^ iBMMs. Data are normalized to WT iBMMs for each strain. (B) Intracellular growth of wildtype *Lm* in WT and *Pten*^−/−^iBMMs. Data are normalized to CFU at 30 minutes for each genotype. (C) Immunofluorescence microscopy of infected WT or *Pten*^−/−^ iBMMs. Cells were infected by centrifugation with mCherry-*Lm* at MOI=5 and fixed 15 min post-infection. Extracellular bacteria were labeled with *Listeria* antisera and a fluorescent secondary antibody. Macrophage nuclei were stained with DAPI. Scale bars indicate 25 μm. (D) Quantification of adherence in (C). The total number of bacteria per 100 macrophages was counted for at least three fields of view (approximately 600 macrophages) per genotype and averaged for each experiment. (E) Quantification of *Lm* internalization 1 hour post-infection. The percentage of intracellular bacteria was calculated for at least three fields of view per genotype and averaged for each experiment. All data are means and SEM of at least three biological replicates. \*\**p*<0.01, \*\*\**p*<0.001, n.s. = non-significant, as determined by unpaired *t* tests (A) or ratio paired *t* tests (D, E).

To further investigate the role of PTEN in *Lm* internalization, we employed an immunofluorescence strategy to quantify intracellular and extracellular bacteria over time during macrophage infection. WT and *Pten*^−/−^ iBMMs were cultured on glass coverslips and infected with mCherry-expressing *Lm* to visualize all adherent and intracellular bacteria. At various timepoints, coverslips were washed to remove non-adhered bacteria, fixed, and immunostained under non-permeabilizing conditions to label only extracellular adherent bacteria. Importantly, gentamicin was not used in these experiments, allowing simultaneous measurement of both adherence and internalization. At 15 minutes post-infection, *Pten*^−/−^ iBMMs had far fewer bacteria present overall (Figure 3C and 3D), indicating a defect in *Lm* adherence to *Pten*^−/−^ macrophages relative to WT iBMMs. To measure efficiency of internalization, we calculated the percentage of total bacteria that were intracellular for each genotype. There was no difference in the percentage of intracellular bacteria between WT and *Pterr*^−/−^ macrophages (Figure 3E), suggesting that PTEN does not affect the internalization rate of *Lm*. We observed similar results over an entire 1 hour time course for both adherence and internalization, with the adherence defect in *Pten*^−/−^ iBMMs apparent as early as 5 minutes post-infection (S3 Fig). Collectively, our data indicate that PTEN enhances phagocytosis by promoting adherence of *Lm* to macrophages.

### The lipid phosphatase activity of PTEN promotes *Lm* uptake

PTEN is a protein and lipid phosphatase which participates in major phospho-signaling cascades in mammalian cells (27). To begin dissecting the PTEN-dependent signaling pathways involved in regulation of *Lm* adherence, we first investigated the enzymatic activity of PTEN required for *Lm* uptake. The following PTEN variants were constitutively expressed in *Pten* iBMMs: a CRISPR-resistant PTEN allele (PTEN^CR^); PTEN^CR^ G129E, which retains protein phosphatase activity but is unable to dephosphorylate lipids; or PTEN^CR^ G129R, which lacks both protein and lipid phosphatase activity (33). Expression of each PTEN variant was confirmed at the protein level by immunoblot (Figure 4A). Each cell line was infected with *Lm* and uptake was measured by gentamicin protection assay. Ectopic expression of PTEN^CR^ in *Pten*^−/−^ iBMMs restored uptake of *Lm* to that of WT iBMMs (Figure 4B). In contrast, neither the lipid phosphatase mutant (G129E) nor the enzymatically dead mutant (G129R) variants of PTEN rescued *Lm* uptake in *Pten* iBMMs, demonstrating that the lipid phosphatase activity of PTEN is required for enhancement of *Lm* phagocytosis.

**Fig 4.**
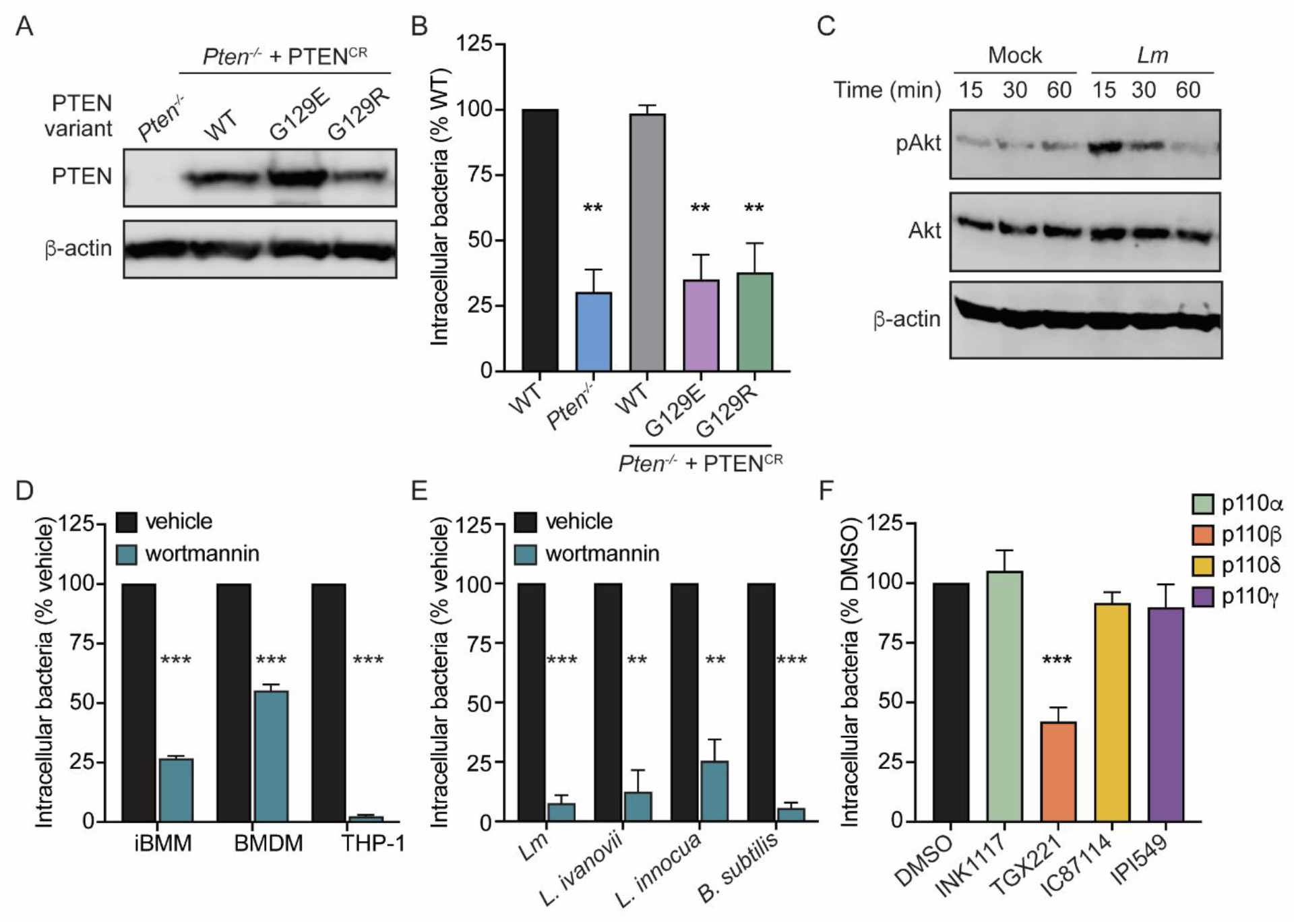
The role of PIP_3_ in regulating phagocytosis of *Lm* by macrophages. (A) Immunoblot of PTEN in *Pten*^−/−^ iBMMs expressing CRISPR-resistant PTEN (PTEN^CR^), PTEN^CR^ G129E, or PTEN^CR^ G129R. β-actin was used as a loading control. (B) Gentamicin protection assay measuring uptake of *Lm* by WT iBMMs or *Pten*^−/−^ iBMMs expressing PTEN variants. Data are normalized to WT iBMMs. (C) Immunoblot of phosphorylated Akt (Ser473) in BMDMs in response to *Lm* infection. BMDMs were mock infected or infected with MOI=100 and lysed at the indicated timepoints. Total Akt and β-actin were probed as loading controls. (D) Gentamicin protection assay measuring uptake of *Lm* by macrophages treated with 100 nM wortmannin. Data are normalized to vehicle-treated cells. (E) Gentamicin protection assay measuring uptake of bacteria by iBMMs in the presence of 100 nM wortmannin. Data are normalized to vehicle-treated cells. (F) Gentamicin protection assay measuring uptake of *Lm* by iBMMs in the presence of 1 μM INK1117, 10 μM TGX221, 10 μM IC87114, or 50 nM IPI549. Data are normalized to DMSO-treated controls. All data are means and SEM of at least three biological replicates. \*\**p*<0.01, \*\*\**p*<0.001, as determined by unpaired *t* tests.

### Class 1A PI3K activation is required for *Lm* phagocytosis by macrophages

The lipid phosphatase activity of PTEN specifically dephosphorylates the D3 position of 3-phosphorylated phosphatidylinositol species, mainly phosphatidylinositol (3,4,5)-triphosphate (PIP_3_) (28). PIP_3_ is transiently produced upon phosphorylation of PI(4,5)P_2_ (PIP_2_) by activated PI3K and turnover of PIP_3_ back to PIP_2_ is mediated by PTEN via dephosphorylation (27). We hypothesized that turnover of PIP_3_ by PTEN is important for enhancement of *Lm* phagocytosis. To begin investigating the role of PIP_3_ in *Lm* uptake, we first determined whether PIP_3_ is produced during *Lm* infection of macrophages. Upon PI3K activation and subsequent PIP_3_ production, Akt is recruited to the plasma membrane where it undergoes phosphorylation (34); therefore, detection of phosphorylated Akt is a common method of measuring PI3K activation. Akt phosphorylation was measured by immunoblot analysis at 15, 30, and 60 minutes post-infection. Infection with *Lm* induced strong Akt phosphorylation 15 minutes post-infection, which waned at 30 minutes and returned to baseline levels by 60 minutes post-infection (Figure 4C). These data indicate that *Lm* rapidly and transiently activates PI3K and PIP_3_ production during macrophage infection.

We next modulated PIP_3_ levels during infection and measured the effect on *Lm* uptake. First, PIP_3_ production was blocked by treating macrophages with the PI3K inhibitor wortmannin. iBMMs, BMDMs, and THP-1 cells all exhibited reduced *Lm* uptake when pre-treated with wortmannin (Figure 4D), indicating that PI3K activation is required for phagocytosis of *Lm* by macrophages. Given that PTEN was specifically required for uptake of *Lm* compared to other bacterial species, we also assessed the specificity of the requirement for PI3K. In contrast to PTEN, PI3K activation was required for uptake of *L. ivanovii, L. innocua*, and *B. subtilis* (Figure 4E), suggesting that PI3K activation is broadly important for phagocytosis of bacteria by macrophages.

In mammals, PI3Ks encompass a large family of enzymes consisting of three classes that phosphorylate different phosphoinositides, all of which are inhibited by wortmannin. The majority of PI3K isoforms belong to Class I, which produces PIP_3_ from PI(4,5)P_2_, while Class II and Class III PI3Ks produce other phosphoinositide species such as PI3P and PI(3,4)P_2_ (35). The catalytic subunits of Class I PI3Ks are further divided into Class 1A isoforms (p110α, p110β, and p110δ) and a single Class 1B isoform (p110γ). To identify whether PIP_3_-producing PI3K isoforms are involved in promoting phagocytosis of *Lm*, we pre-treated iBMMs with isoform-specific PI3K inhibitors targeting all four Class I PI3K isoforms and measured uptake of *Lm*. Inhibition of p110β blocked uptake of *Lm* in iBMMs, while inhibition of p110α, p110δ, or p110γ had no effect (Figure 4F). Collectively, these data show that PIP_3_ production by the Class 1A PI3K p110β is required for phagocytosis of *Lm* by macrophages.

Class 1A PI3Ks are canonically activated by phosphorylated tyrosine residues on receptor tyrosine kinases and other cell surface receptors (35). Additionally, Class 1A PI3Ks are activated downstream of toll-like receptors (TLRs) in response to bacterial infection (36,37). Given that *Lm* is known to activate TLR2 and TLR5 on the surface of macrophages (38,39), we hypothesized that *Lm*-induced PI3K activation in macrophages is mediated by TLRs. To test this hypothesis, we measured uptake of *Lm* and PI3K activation in *Tlr2*^−/−^ macrophages. For these studies, BMDMs from *Tlr2*^−/−^ mice were isolated and absence of TLR2 was confirmed by immunoblot analysis (S4A Fig). Interestingly, uptake of *Lm* was unchanged in *Tlr2*^−/−^ BMDMs compared to WT BMDMs (S4B Fig). While we did not directly test *Tlr5*^−/−^ macrophages, studies from our lab have found that *Lm* lacking the TLR5 ligand *flaA* are taken up by macrophages with equal efficiency as wildtype *Lm* (40). To account for the possibility of redundancy between TLR2 and TLR5, we infected wildtype and *Tlr2*^−/−^ BMDMs with Δ*flaA* and found that uptake remained unchanged in the absence of TLR2 and TLR5 recognition (S4C Fig). These results indicate that phagocytosis of *Lm* is TLR-independent.

To test the role of TLRs in mediating PI3K/PTEN signaling in macrophages during *Lm* infection, we pre-treated WT and *Tlr2*^−/−^ BMDMs with PI3K and PTEN inhibitors prior to infection with Δ*flaA*. Wortmannin and bpV(pic) inhibited uptake of Δ*flaA* to a similar extent in both WT and *Tlr2*^−/−^ BMDMs (S4D Fig), suggesting that PI3K signaling is intact in the absence of TLR stimulation. To test this, we monitored Akt phosphorylation in WT and *Tlr2*^−/−^ BMDMs infected with wildtype *Lm* and Δ*flaA* and observed Akt phosphorylation over background in the absence of TLR2 and TLR5 recognition (S4E Fig). Together, these data demonstrate that *Lm*-induced PI3K activation in macrophages occurs independently of TLR2 and TLR5. Given that specific *Lm* strains are phagocytosed using a distinct PTEN-dependent pathway compared to other *Lm* strains or *L. innocua*, we tested whether PTEN-independent uptake of the latter was mediated by TLRs. To that end, we compared uptake of *Lm* 10403S, *Lm* Li2, and *L. innocua* in WT and *Tlr2*^−/−^ BMDMs and found that uptake of all three bacterial strains was TLR2-independent (S4F Fig). While we did not directly test a role for TLR5 in PTEN-independent uptake, *Lm* Li2 and *L. innocua* do not produce flagella at 37°C (41,42), strongly suggesting that these strains do not engage TLR5 in our assays. Overall, our data show that uptake of *Listeria* species in murine macrophages is TLR-independent.

### Myeloid PTEN promotes bacterial restriction during murine foodborne listeriosis

To investigate the role of PTEN-dependent phagocytosis of *Lm in vivo*, we generated myeloid-specific conditional *Pten* knockout mice. C57BL/6 mice expressing Cre recombinase from the myeloid-specific *Lyz2* promoter (*LysM-Cre^+/+^*) were crossed with C57BL/6 mice containing *loxP* sites flanking exon 5 of *Pten* (*Pten*^fl/fl^ to produce *LysM*-Cre^+/+^ *Pter*^fl/fl^ myeloid knockout mice (PTEN^M-KO^). *LysM*-Cre^+/+^ *Pten^wt/wt^* mice which express Cre but do not contain *Pten loxP* sites (PTEN^M-WT^) were included as controls. We used a foodborne model of listeriosis to ensure passage through the intestinal lamina propria where *Lm* is engulfed by resident macrophages, which are hypothesized to promote dissemination of *Lm* to peripheral organs (3,4). 8-10 week old PTEN^M-WT^ and PTEN^M-KO^ sex- and age-matched mice were given streptomycin in their drinking water for 48 hours and fasted for 16 hours prior to infection to increase susceptibility to oral infection (43,44). Mice were then fed 10^8^ *Lm* in PBS and body weight was recorded over the course of 5 days as an overall measurement of disease severity. To assess bacterial burdens throughout the infection, mice were euthanized at 1, 3, and 5 days post-infection (dpi) and CFU were enumerated from organs. In addition, ilea and colons were fractionated to separate the mucus layer, epithelial cells, and lamina propria compartments and CFU were enumerated from each fraction.

At 1 dpi, we observed increased bacterial burdens in the epithelial cells and lamina propria of the ileum in PTEN^M-KO^ mice compared to PTEN^M-WT^ mice (Figure 5A). There was no difference in CFU in the ileum mucus layer (Figure 5A), suggesting that higher bacterial burdens in the epithelial cells and lamina propria were not due to increased colonization of the ileal lumen. In contrast to the lamina propria, there were no differences in CFU between PTEN^M-WT^ and PTEN^M-KO^ mice in the mesenteric lymph nodes (MLN), spleen, or liver 1 dpi (Figure 6 D-F), indicating that myeloid PTEN is dispensable for initial dissemination of *Lm* to peripheral organs. We also observed no difference in CFU in the feces or cecum between PTEN^M-WT^ and PTEN^M-KO^ mice throughout the experiment (Figure 6B and 6C), demonstrating that PTEN does not affect colonization of the GI tract or fecal shedding. Together, these data show that myeloid expression of PTEN restricts *Lm* replication in the epithelial cells and lamina propria of the ileum early during infection, however increased bacterial burdens in these compartments does not affect dissemination.

**Fig 5.**
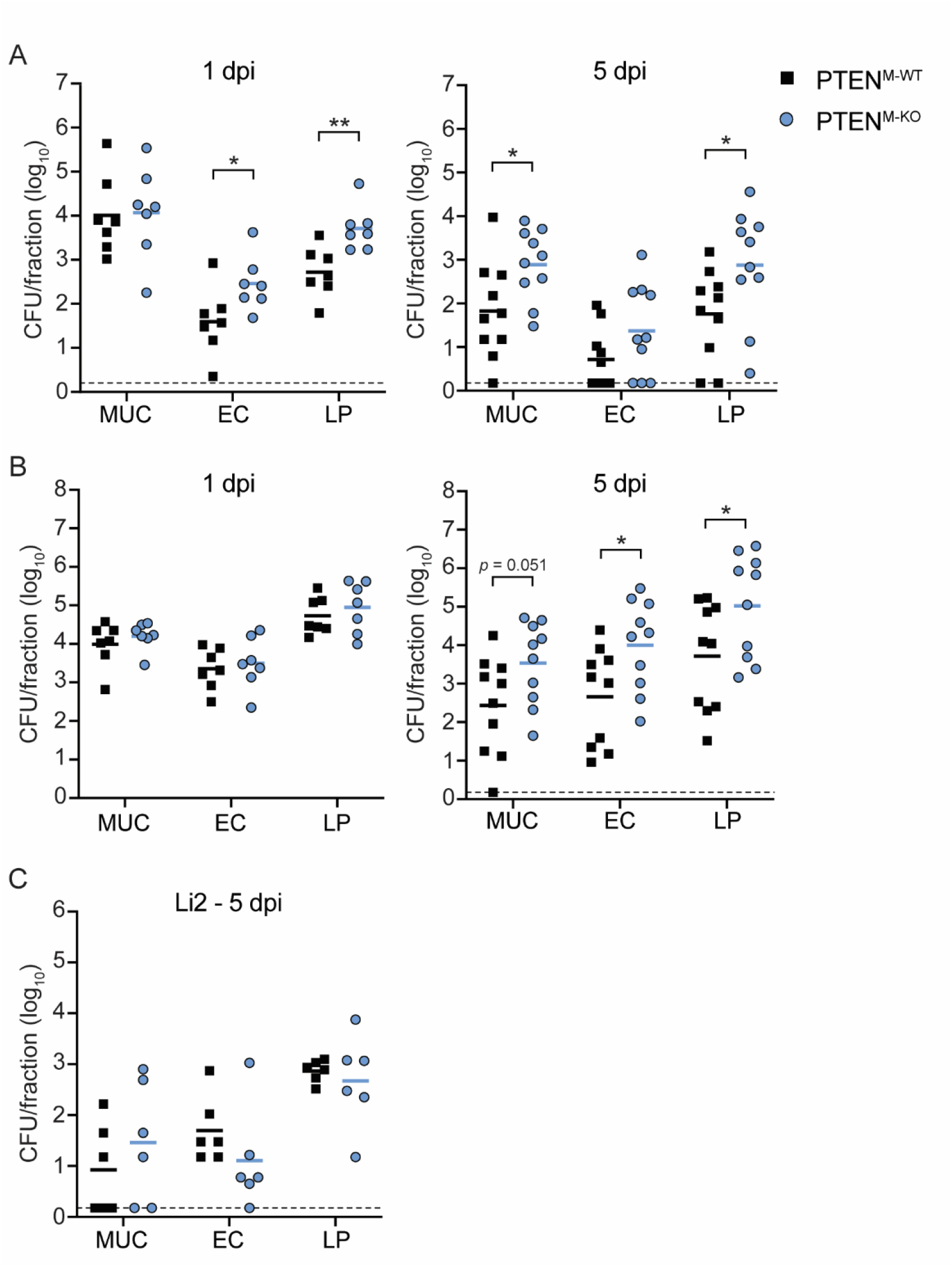
Myeloid PTEN restricts bacterial replication in the intestinal lamina propria. Mice were orally infected with 10^8^ *Lm* 10403S or 5 x 10^8^ *Lm* Li2. Bacterial burdens in (A) ileal and (B) colonic intestinal fractions were enumerated 1 and 5 dpi (MUC = mucus layer, EC = epithelial cells, LP = lamina propria). Each data point represents a single mouse (1 dpi, n = 7 per genotype; 5 dpi, n = 10 per genotype for 10403S or n = 6 per genotype for Li2). Data from 10403S-infected animals represent two independent experiments per timepoint. Solid lines indicate geometric means. Dashed lines indicate the limit of detection (l.o.d.). \**p*<0.05, \*\**p*<0.01 as determined by unpaired *t* tests of natural log-transformed values.

**Fig 6.**
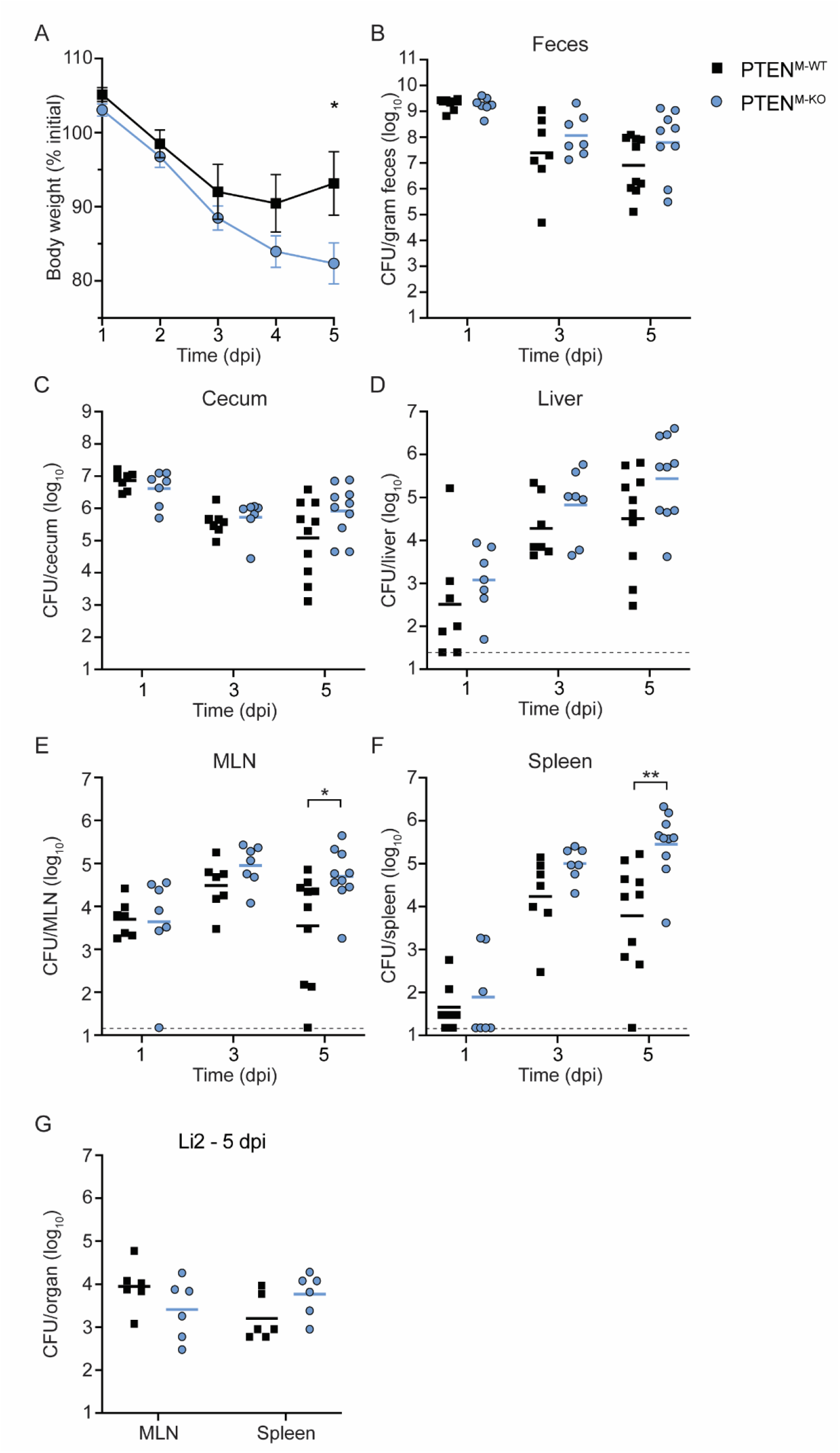
Myeloid PTEN promotes bacterial clearance during murine foodborne listeriosis. (A) Body weights of infected mice over time, reported as the percent of initial weight before streptomycin treatment. Data are means and SEM of n = 10 mice at each timepoint. (B-F) Mice were orally infected with 10^8^ *Lm* 10403S and CFU from tissues were enumerated at 1, 3, and 5 dpi. Each data point represents a single mouse (1 and 3 dpi, n = 7 per genotype; 5 dpi, n = 10 per genotype). Data for each time point represents two independent experiments. Solid lines indicate geometric means. Dashed lines represent the l.o.d.. (**G**) Mice were orally infected with 5 x 10^8^ *Lm* Li2 and CFU from tissues were enumerated 5 dpi. Each data point represents a single mouse (n = 6 per genotype). Solid lines indicate geometric means. \**p*<0.05, \*\**p*<0.01 as determined by unpaired *t* tests of untransformed (A) or natural log-transformed (B-G) values.

There were no differences in weight loss or bacterial burdens in any organs tested at 3 dpi (Figures 6A-F and S5 Fig). By 5 dpi, however, PTEN^M-WT^ mice lost ~7% of their initial body weight while PTEN^M-KO^ mice lost nearly 18% of their initial weight (Figure 6A), indicating that PTEN^M-KO^ mice experienced more severe disease during the later stages of infection. Consistent with increased weight loss, PTEN^M-KO^ mice displayed increased bacterial burdens in the MLN (~1-log) and spleen (~2-logs) compared to PTEN^M-WT^ mice at 5 dpi (Figure 6E and 6F). In addition, the lamina propria of both the ileum and colon as well as the ileal mucus layer and colonic epithelium of PTEN^M-KO^ mice also contained higher *Lm* burdens compared to PTEN^M-WT^ mice at 5 dpi (Figure 5A and 5B), suggesting that PTEN^M-KO^ mice contained higher bacterial burdens overall in these organs. Taken together, these data demonstrate that myeloid PTEN limits disease severity and restricts *Lm* growth during later stages of foodborne listeriosis.

To specifically assess the contribution of PTEN-regulated phagocytosis to controlling *Lm* infection, we infected mice with the *Lm* strain Li2, which is phagocytosed using a PTEN-independent mechanism (Figure 2H). We reasoned that any global effects of myeloid PTEN loss would impact both strains equally, while phagocytosis-specific phenotypes would not affect *Lm* Li2 infection. In contrast to mice infected with 10403S, there were no significant differences in CFU of any organs between PTEN^M-WT^ and PTEN^M-KO^ mice infected with Li2 5 dpi, including the MLN, spleen, and colon fractions (Figures 5B and 6G, S6 Fig). Collectively, these data suggest that restriction of *Lm* 10403S by myeloid PTEN is driven by PTEN-dependent phagocytosis.

## Discussion

Host determinants of opsonin-independent phagocytosis of *Lm* have not previously been studied. Here, we performed a genome-wide CRISPR/Cas9 screen that identified over 200 macrophage genes necessary for *Lm* infection, the vast majority of which have not previously been implicated in *Lm* pathogenesis. Among the candidate genes identified by the screen were several members of the PTEN signaling pathway, including PTEN itself. PTEN promoted phagocytosis of *Lm* by enhancing adherence of the bacteria to macrophages. Both PIP_3_ production by the type 1A PI3K p110β and the lipid phosphatase activity of PTEN are required for phagocytosis, strongly suggesting that PIP_3_ turnover is important for enhancement of *Lm* uptake. The role of PTEN-dependent phagocytosis *in vivo* was assessed using an oral *Lm* infection of myeloid conditional knockout mice. Myeloid PTEN limits bacterial burdens in the lamina propria early during infection, however this does not impact initial dissemination to internal organs. During later stages of infection, myeloid PTEN promotes bacterial restriction in several organs and protects against severe disease. Taken together, our results indicate a novel role for PTEN in regulating phagocytosis of *Lm* and begin to characterize the contributions of opsonin-independent phagocytosis *in vivo*.

Our results demonstrate that PTEN promotes phagocytosis of *Lm* specifically by macrophages. These data are surprising and intriguing for a few reasons. First, it is well-documented that class I PI3Ks are required for the phagocytic activity of macrophages due to PIP_3_ regulation of actin polymerization at the phagocytic cup (45). Consistent with this, we and others have found that phagocytosis of several bacterial species is dependent on PI3K activation (46–49), suggesting that PI3K activation and PIP_3_ production are common features of bacterial recognition by macrophages. PTEN has thus been characterized as a repressor of phagocytosis due to PIP_3_ dephosphorylation and antagonism of PI3K signaling (50–53). To our knowledge, this is the first report that PTEN can promote phagocytosis of bacteria by macrophages. In addition, PTEN appears to play opposing roles in regulating uptake of *Lm* by phagocytes and non-phagocytes. PI3K activation is also required for internalin-mediated uptake of *Lm* into non-phagocytic cells (54), suggesting that PTEN would act as a repressor of this process. Indeed, we found that inhibition of PTEN increased uptake of *Lm* by hepatocytes and intestinal epithelial cells. Our finding that PTEN promotes phagocytosis of *Lm* by enhancing adherence to macrophages provides a possible explanation for these cell type differences, whereby PTEN regulates macrophage-specific *Lm* adherence factors. Overall, our results support a model in which PI3K activation and PIP_3_ production are broadly required for internalization of bacteria, including *Lm*, and PTEN functions distinctly to promote adherence of *Lm* to macrophages.

The factors mediating adherence of *Lm* to macrophages are not well understood. We hypothesize that PTEN affects adherence to human and murine macrophages by increasing availability of one or more surface receptors which bind *Lm*. Elevated baseline PIP_3_ levels in *Pten*^−/−^ cells may serve in a signaling capacity to regulate expression of *Lm*-specific receptors or altered membrane dynamics may affect surface presentation of adherence factors. Alternatively, turnover of PIP_3_ by PTEN may be important for regeneration of PI(4,5)P_2_, which is capable of acting in a signaling capacity to regulate membrane dynamics (55,56). Our data indicate that TLRs do not serve as primary phagocytic receptors for *Lm*, as phagocytosis of *Lm* and *L. innocua* were TLR-independent. We demonstrated that the class 1A PI3K p110β was specifically required for phagocytosis of *Lm*, and p110β is canonically activated by receptor tyrosine kinases (RTKs) (35), suggesting that phagocytosis of *Lm* may involve one or more RTKs. Very few macrophage receptors have been previously implicated in phagocytosis of non-opsonized *Lm*. Fcγ-receptor 1A (FcγR1a or CD64) mediates antibody-independent uptake of *Lm* specifically by human macrophages, although murine FcγR1a had no effect on internalization of *Lm* (57). The macrophage scavenger receptor SR-A I/II (*Msr1*) reportedly promotes phagocytosis of *Lm* by peritoneal macrophages and Kupffer cells during i.v. infection of mice, however internalization via this pathway is listericidal (58). The identity of *Lm*-specific receptor(s) which activate PI3K and lead to cytosolic *Lm* replication in both human and mouse macrophages remains to be determined and is an active area of investigation.

The finding that different strains of *Lm* undergo distinct mechanisms of uptake is intriguing and indicates that both host and bacterial factors regulate phagocytosis of *Lm*. Interestingly, the strains of *Lm* internalized via PTEN-dependent phagocytosis belong to serovar 1/2, while strains whose uptake is PTEN-independent belong to serovar 4 (S2 Table). The 16 serovars of *Lm* are differentiated by the chemical composition of the cell wall-associated teichoic acids (59). Thus, while this is not an exhaustive characterization of *Lm* serovars, our data suggest that differences in bacterial surface molecules dictate whether *Lm* is phagocytosed primarily via a PTEN-dependent or -independent mechanism, likely through recognition by distinct receptors. *Lm* serovar is associated with disease severity, with the majority of human listeriosis cases caused by serovar 4b strains (2). Recently, specific modifications of wall teichoic acids on serovar 4b strains were found to be important for pathogenesis by promoting invasion of non-phagocytic cells (60). Given that PTEN-dependent phagocytosis is a host defense against listeriosis, as discussed below, it is possible that avoidance of phagocytosis through this pathway by serovar 4b strains may contribute to their virulence.

To dissect the role of PTEN-dependent phagocytosis *in vivo*, we infected myeloid PTEN conditional knockout mice (PTEN^M-KO^) via the oral inoculation route. This approach gave us a unique opportunity to study the contribution of macrophage phagocytosis to *Lm* disease progression during foodborne listeriosis. In the absence of myeloid PTEN, bacterial burdens were elevated in the ileal lamina propria early during infection. These results were surprising, as macrophages are thought to be the main growth-permissive cell type in the lamina propria, leading us to predict reduced bacterial burdens in PTEN^M-KO^ mice as a result of reduced uptake into macrophages. It was recently demonstrated that infection of specific macrophage subsets in the lamina propria is important for early production of IFNγ and promotes bacterial restriction in this compartment (61). It is therefore possible that reduced macrophage phagocytosis in PTEN^M-KO^ mice blunts local immune responses that normally restrict replication of *Lm* in the lamina propria. Nonetheless, the localization of *Lm* in the lamina propria of PTEN^M-KO^ mice is currently unknown. The majority of bacteria in the lamina propria are extracellular following oral infection (3), suggesting that *Lm* may be able to more efficiently replicate in the extracellular space in the absence of macrophage phagocytosis. Alternatively, lymphocytes support intracellular growth of *Lm in vivo* (62), and may serve as a replicative niche in the lamina propria of PTEN^M-KO^ mice. *Lm* does not replicate in dendritic cells or monocytes in the gut (63,64), so these populations are unlikely to serve as a replication niche. While the exact cell types responsible for promoting *Lm* growth in and dissemination from the lamina propria are not well understood, our data suggest that macrophages play a primarily protective role in the gut.

Mice lacking myeloid PTEN had higher bacterial burdens in the intestinal tissue, MLN, and spleen during later stages of infection. Importantly, these differences were not observed during infection of the *Lm* serovar 4b strain Li2, indicating that this phenotype is driven by PTEN-regulated phagocytosis rather than global effects of myeloid PTEN loss. Overall, our *in vivo* studies demonstrate that macrophage phagocytosis is important for growth restriction and clearance of *Lm* during foodborne listeriosis. These data are consistent with the demonstrated role of macrophages in host protection during i.v. infection of mice, in which IFNγ-primed macrophages are critical (14,15). Recent work has demonstrated a similar requirement of IFNγ for bacterial restriction 4 days after oral infection with *Lm* (61). We hypothesize that IFNγ-activated macrophages are unable to efficiently phagocytose and clear *Lm* in PTEN^M-KO^ mice, leading to increased bacterial burdens. Alternatively, reduced uptake by macrophages early during infection may dampen cytokine production by these cells, which are important for establishing a robust immune response. Although few studies have investigated the immune response to *Lm* during oral infection (65), development of foodborne infection models of murine listeriosis has paved the way for studying interactions between *Lm* and immune cells in a physiologically relevant system (66). Our infection studies have contributed to this body of work by demonstrating the protective role of macrophages during oral infection and providing a unique murine model to study macrophage phagocytosis of *Lm in vivo*.

Taken together, the genome-wide CRISPR screen described here identified hundreds of novel candidate genes important for macrophage infection by *Lm*. These data provide a valuable resource to investigate *Lm* pathogenesis and phagocytosis of bacteria more broadly. In contrast to the current understanding of bacterial phagocytosis as a process driven by recognition of conserved PAMPs (67,68), our work demonstrates that highly related bacteria are internalized via functionally distinct mechanisms, even down to the strain level. Finally, we have defined a role for macrophages as an important host defense during foodborne listeriosis and provide the first evidence that non-opsonic phagocytosis contributes to *Lm* pathogenesis.

---

**S1 Fig.**
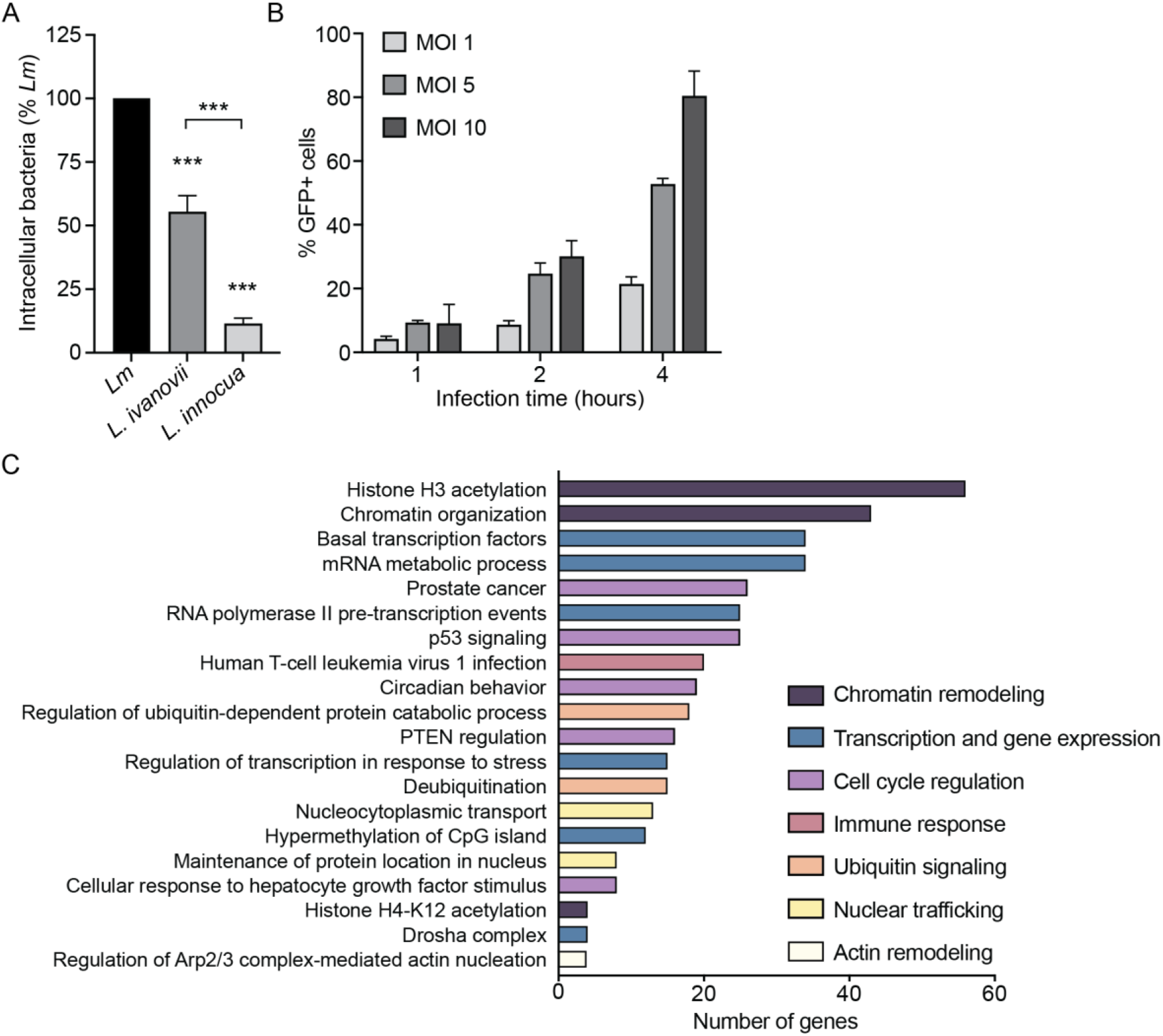
CRISPR screen optimization and Metascape analysis. (A) Gentamicin protection assay measuring bacterial uptake in iBMMs expressing Cas9. Data are normalized to uptake of *Lm*. (B) Optimization of infection efficiency. iBMMs were infected with GFP-*Lm* for the indicated MOI and time, and GFP^+^ cells were quantified by flow cytometry 6 hours post-infection. (C) Hits with *p*<0.01 (n=235) were analyzed by Metascape. The number of genes belonging to each biological process is depicted. Colors indicate pathways with similar functions. Data in (A-B) are means and SEM of three biological replicates. \*\*\**p*<0.001 as determined by unpaired *t* tests.

**S2 Fig.**
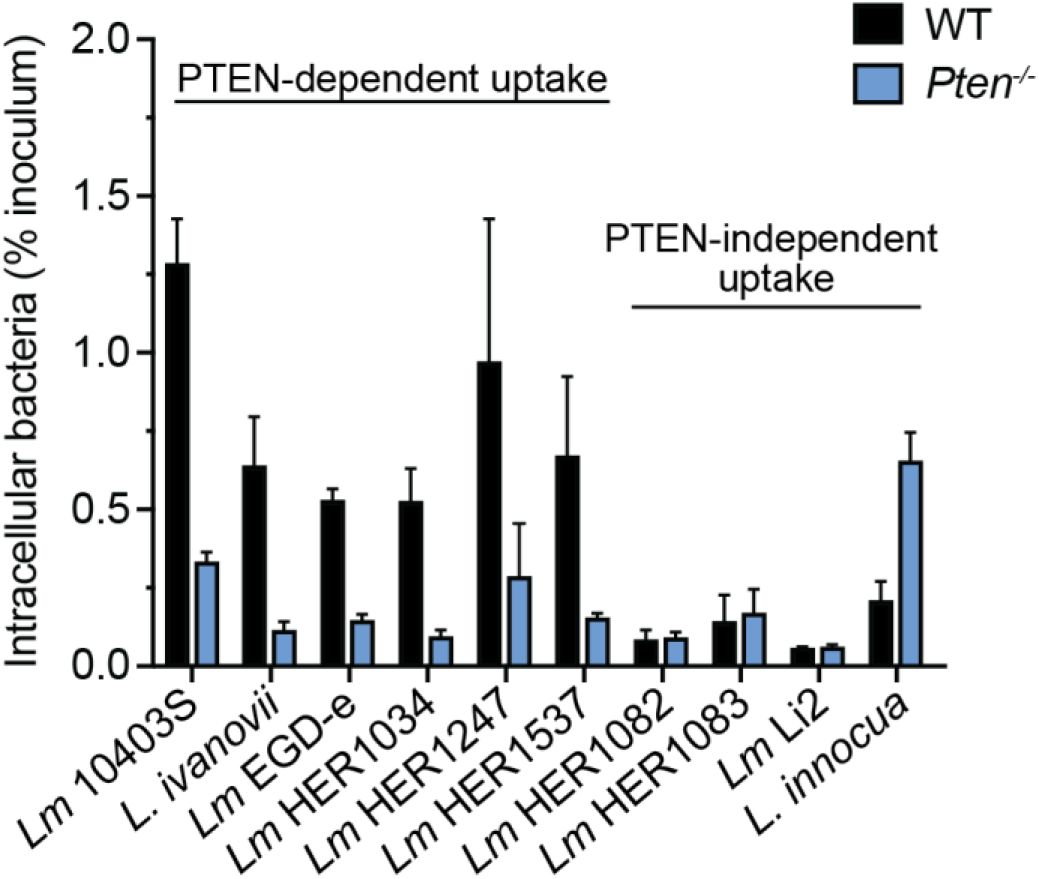
Enhanced phagocytosis of *Lm* is PTEN-dependent. Gentamicin protection assay measuring uptake of *Lm* strains, *L. ivanovii*, and *L. innocua* by iBMMs. The initial inoculum of each strain was enumerated and the percentage of internalized bacteria was calculated. Data are means and SEM of at least two biological replicates.

**S3 Fig.**
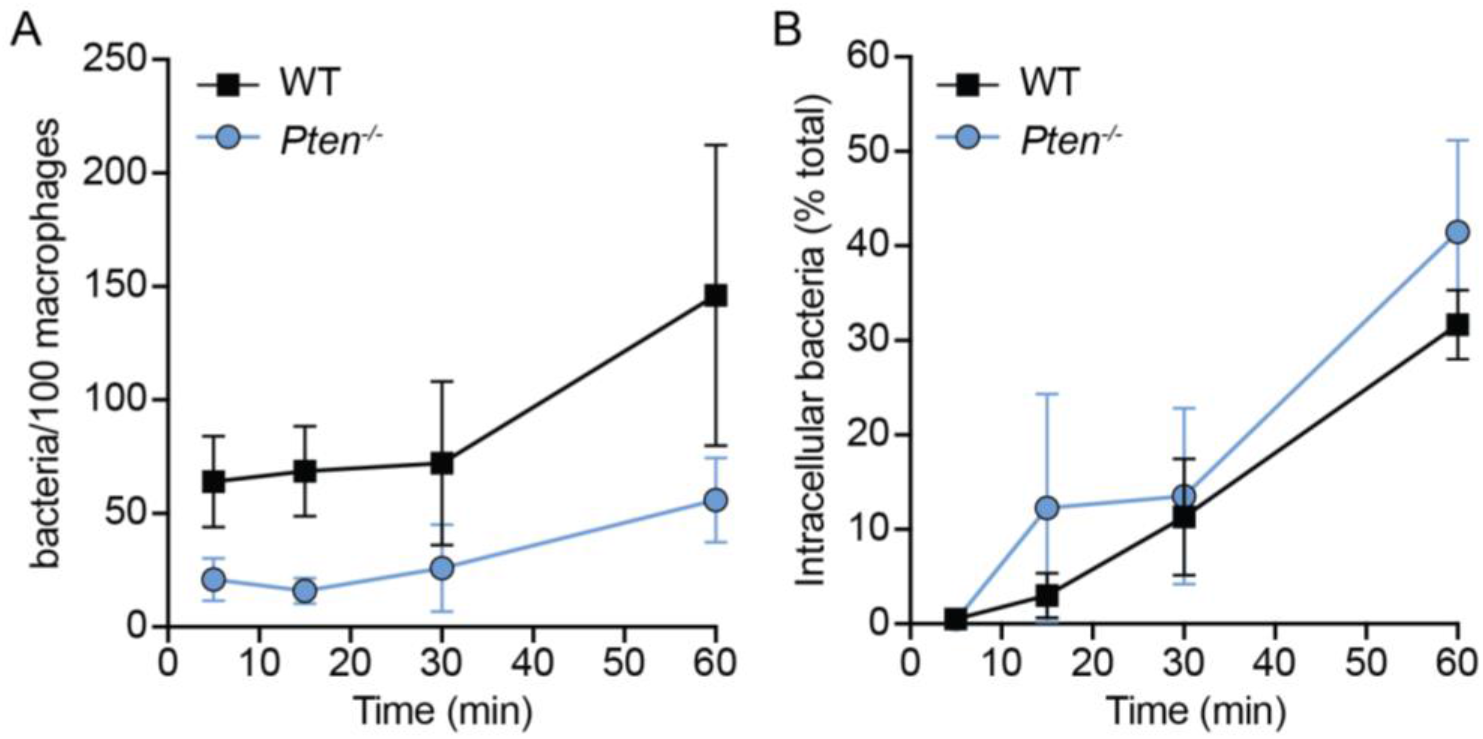
Dynamics of *Pten*^−/−^ iBMM infection. iBMMs were infected, immunostained, and quantified as in Figure 3C-E. Timepoints were taken at 5, 15, 30, and 60 minutes post-infection. (**A**) Adherence and (**B**) internalization of *Lm* by iBMMs during the 1 hour time course were quantified. All data are means and SEM of three biological replicates.

**S4 Fig.**
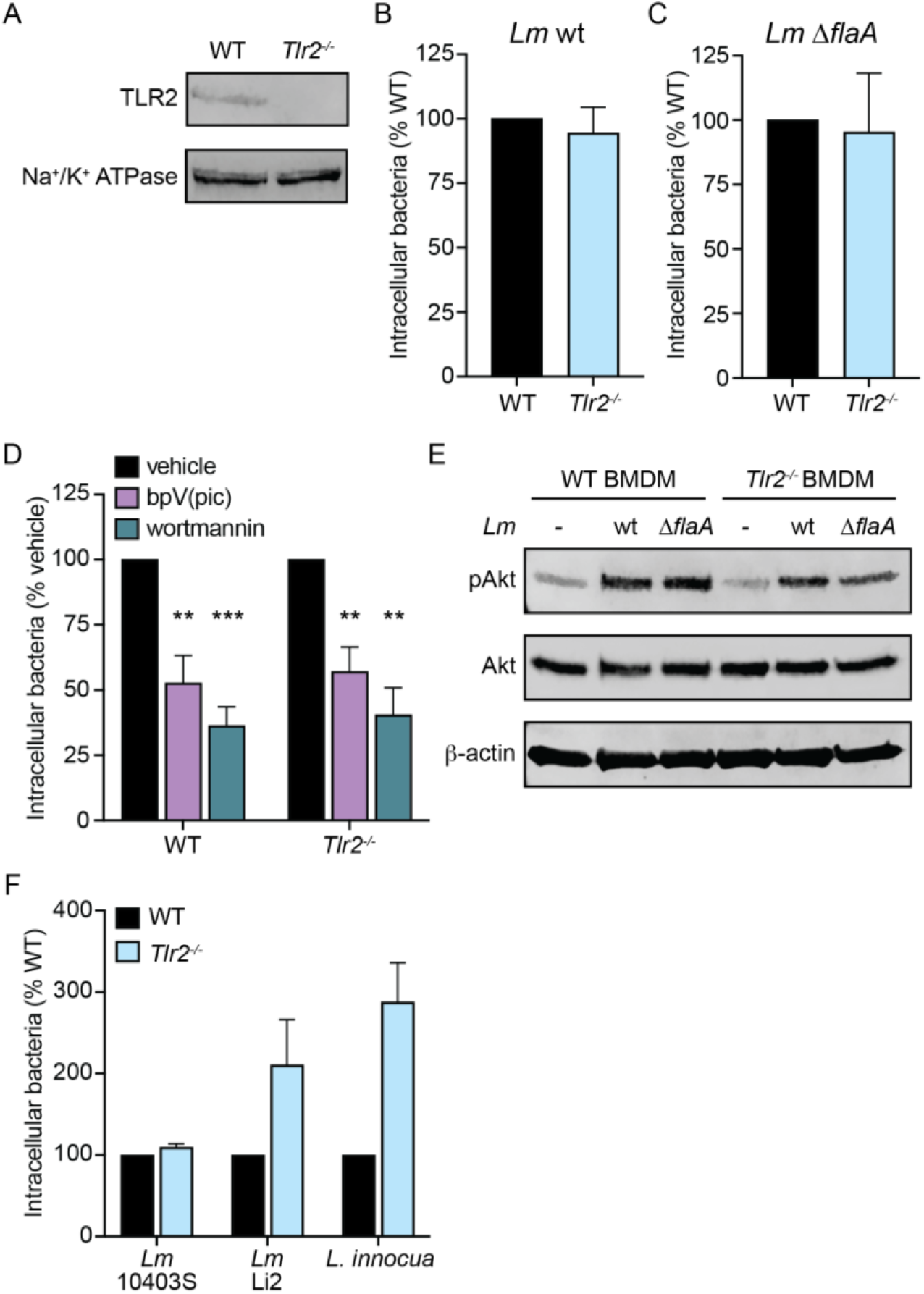
Uptake of *Listeria* by BMDMs is TLR-independent. (**A**) Immunoblot analysis of TLR2 in WT and *Tlr2*^−/−^ BMDMs. Na^+^/K^+^ ATPase was used as a loading control for membrane proteins. (**B**) Gentamicin protection assay measuring uptake of wildtype (wt) *Lm* by WT and *Tlr2*^−/−^ BMDMs. Data are normalized to WT BMDMs. (**C**) Gentamicin protection assay measuring uptake of *Lm* Δ*flaA* by WT and *Tlr2*^−/−^ BMDMs. Data are normalized to WT BMDMs. (**D**) Gentamicin protection assay measuring uptake of *Lm ΔflaA* by WT and *Tlr2*^−/−^ BMDMs in the presence of 5 μM bpV(pic) or 100 nM wortmannin. Data are normalized to vehicle-treated cells. (**E**) Immunoblot of phosphorylated Akt (Ser473) in WT and *Tlr2*^−/−^ BMDMs mock-infected or infected with *Lm* wt or Δ*flaA* strains. BMDMs were infected with MOI=100 and lysed 15 minutes post-infection. Total Akt and β-actin were used as loading controls. (**F**) Gentamicin protection assay measuring uptake of *Lm* 10403S, *Lm* Li2, or *L. innocua* by WT and *Tlr2*^−/−^ BMDMs. Data are normalized to WT BMDMs for each strain. All data are means and SEM of at least three biological replicates except (F) which consists of two biological replicates. \*\**p*<0.01, \*\*\**p*<0.001, as determined by unpaired *t* tests.

**S5 Fig.**
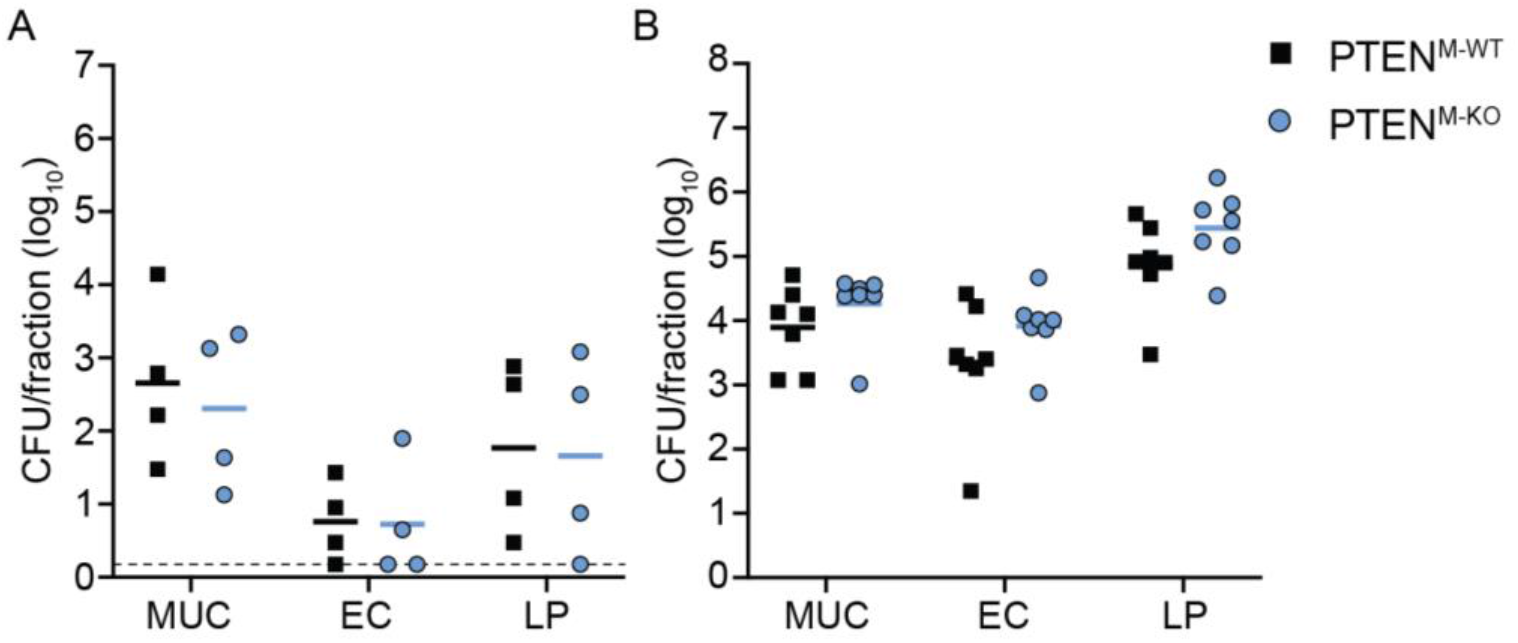
Intestinal fractions from 10403S-infected mice 3 dpi. Mice were orally infected with 10^8^ *Lm* 10403S. Bacterial burdens in (**A**) ileal and (**B**) colonic intestinal fractions were enumerated 3 dpi (MUC = mucus layer, EC = epithelial cells, LP = lamina propria). Each data point represents a single mouse (ileum, n = 4 per genotype; colon, n = 7 per genotype). Data for colonic fractions represent two independent experiments. Solid lines indicate geometric means. Dashed lines indicate the l.o.d.

**S6 Fig.**
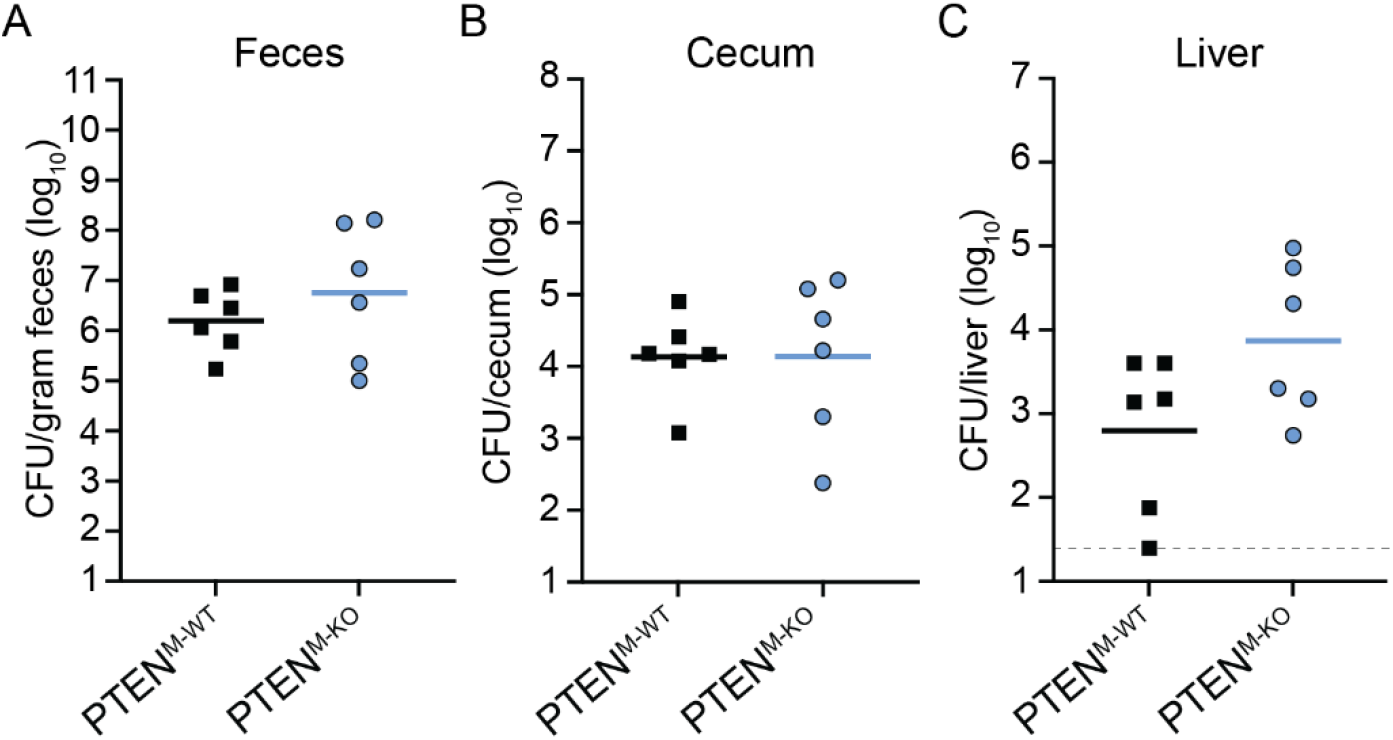
Murine infection with *Lm* Li2 5 dpi. Mice were orally infected with 5 x 10^8^ *Lm* Li2 and CFU were enumerated from (**A**) feces, (**B**) cecum, and (**C**) liver 5 dpi. Each data point represents a single mouse (n = 6 per genotype). Solid lines indicate geometric means. Dashed line indicates the l.o.d.

## Acknowledgements

We would like to thank all members of the Reniere and Woodward Labs at the University of Washington (UW) for valuable feedback and discussions; the Mougous and Lagunoff Labs at UW for reagents and equipment; Dr. Alex Meeske for the gift of several *Listeria* species and *Lm* strains; Dr. Shivam Zaver for critical reading of the manuscript; Dr. Monica Cesinger for assistance with experiments; Serina Tsang at UW In Vivo Services and Dr. Cortney Halsey for assistance with mouse breeding and techniques; and Andy Marty at Fred Hutch Genomics Resource for assistance with Illumina sequencing. This work was supported by NIH research grant R01 AI132356 to M.L.R., R35 GM146795 to A.J.O., and NIH training grants T32AI055396 and T32GM007270 to R.C.G. and M.E.S., respectively.

## Author Contributions

Conceptualization, R.C.G. and M.L.R.; Methodology, R.C.G. and M.L.R.; Formal Analysis, R.C.G. and M.L.R.; Investigation, R.C.G., N.H.S., S.L., M.E.S., M.K.T., and M.L.R.; Resources, A.J.O. and M.L.R.; Writing – Original Draft, R.C.G. and M.L.R.; Writing – Review & Editing, R.C.G., N.H.S., S.L., M.E.S., M.K.T., A.J.O., and M.L.R.; Visualization, R.C.G. and M.L.R.; Funding Acquisition, R.C.G., A.J.O., and M.L.R.

## Materials & Methods

### Materials & data availability

Further information and requests for resources and reagents, including newly generated bacterial strains, cell lines, and plasmids, should be directed to and will be fulfilled by the lead contact, Michelle Reniere. Raw sequencing data from the genome-wide CRISPR screen will deposited at NCBI Sequence Read Archive and be publicly available as of the date of publication. Microscopy data reported in this paper will be shared by the lead contact upon request. Any additional information required to reanalyze the data reported in this paper is available from the lead contact upon request.

### Quantification and statistical Analysis

Statistical analyses were performed using GraphPad Prism 9 and Microsoft Excel. Details on the statistical tests used, number of replicates, and definition of center and error are found in the figure legends. Results with *p*<0.05 are considered statistically significant. Statistical outliers were not omitted in this study.

### Bacterial strains and culture conditions

All bacterial strains used in this study are summarized in S2 Table. Bacteria were cultured on brain heart infusion (BHI) agar (*Listeria spp*.) or LB agar (*B. subtilis, E. coli*) at 37°C. Unless otherwise stated, bacteria were grown in BHI broth overnight at 37°C with shaking prior to experiments. Antibiotics were used at the following concentrations: streptomycin, 200 μg/mL; chloramphenicol, 10 μg/mL (*E. coli*) and 7.5 μg/mL (*Lm*); tetracycline, 2 μg/mL; and carbenicillin, 100 μg/mL. Plasmids were transformed into chemically competent *E. coli* using heat shock.

### Mammalian cell lines

Cas9-expressing immortalized bone marrow-derived macrophages (iBMMs) were described previously (22). THP-1, TIB73, and Caco-2 cell lines were obtained from Dr. Joshua Woodward (University of Washington) (69). HEK293T cells were purchased from ATCC and obtained from Dr. Jennifer Hyde (University of Washington). Cell lines were cultured at 37°C and 5% CO_2_. iBMMs, TIB73, and HEK293T cells were grown in high glucose (4.5 g/L) Dulbecco’s modified Eagle’s medium (DMEM) supplemented with 10% heat-inactivated fetal bovine serum (FBS) (Cytiva), 1 mM sodium pyruvate, 2 mM L-glutamine, and 100 U/mL penicillin-streptomycin (D-10). Caco-2 cells were cultured high glucose DMEM supplemented with 20% FBS, 1 mM sodium pyruvate, 2 mM L-glutamine, and 100 U/mL penicillin-streptomycin (D-20). THP-1 cells were grown in RPMI 1640 supplemented with 10% FBS, 1 mM sodium pyruvate, 2 mM L-glutamine, and 100 U/mL penicillin-streptomycin (RP-10). Cells were plated in medium without penicillin-streptomycin for infections. Cells were determined to be Mycoplasma-free using PCR (70). Antibiotics were used at the following concentrations: gentamicin, 50 μg/mL; puromycin, 5 μg/mL; blasticidin, 10 μg/mL; and Geneticin (G418), 1000 μg/mL.

### Primary cells

Bone marrow-derived macrophages (BMDMs) were cultured as above and grown in high glucose DMEM supplemented with 20% heat-inactivated FBS, 1 mM sodium pyruvate, 2 mM L-glutamine, 10% 3T3 cell supernatant (from M-CSF-producing 3T3 cells), and 55 μM β-mercaptoethanol (BMDM medium). BMDMs were isolated as previously described (71). Briefly, femurs and tibias from 6-8 week old female C57BL/6 mice were crushed with a mortar and pestle in 20 mL BMDM medium and strained through 70 μm cell strainers. Cells were plated in 150 mm untreated culture dishes, supplemented with fresh BMDM medium at day 3, and harvested by pipetting in cold PBS at day 7. BMDMs were aliquoted in 80% BMDM medium, 10% FBS, and 10% DMSO and stored in liquid nitrogen.

### Mice

Animal experiments were carried out in strict accordance with the recommendations in the Guide for the Care and Use of Laboratory Animals of the National Institutes of Health. All protocols were reviewed and approved by the Institutional Animal Care and Use Committee at the University of Washington (Protocol 4410–01). Animals were group-housed in University of Washington Department of Comparative Medicine facilities under specific pathogen-free conditions. 6 week old wildtype (#000664) and *Tlr2*^−/−^ (#004650) female C57BL/6J mice were purchased from The Jackson Laboratory for isolating BMDMs. To generate *Pten* conditional knockout mice, male *LysM*-Cre^+/+^ mice (#004781) and female *Pten*^fl/fl^ mice (#006440) were purchased from The Jackson Laboratory and bred to produce double heterozygotes. Female heterozygotes were bred back to male *LysM*-Cre^+/+^ mice to generate *LysM*-Cre^+/+^ *Pten*^fl/wt^ breeders, which were then bred together to produce *LysM*-Cre^+/+^ *Pten*^fl/fl^ knockout mice (PTEN^M-KO^) and *LysM*-Cre^+/+^ *Pten*^wt/wt^ controls (PTEN^M-WT^). Genotyping was performed by isolating gDNA from ear biopsies and following PCR protocols provided by The Jackson Laboratory. 8-10 week old male and female mice were used for infection studies. Animals were assigned to experimental groups by age-, sex-, and littermate-matching and groups were co-housed when possible.

### Bacterial strain construction

For the CRISPR/Cas9 screen, we constructed a Δ*actA* strain expressing GFP constitutively and RFP under control of the *actA* promoter. RFP expression was used for experiments not described in this manuscript. GFP driven by the constitutive pHyper promoter was cloned into the integrative vector pPL1 (72) by restriction cloning (pPL1.pHyper-GFP). RFP was cloned downstream of the *actA* promoter into the integrative plasmid pPL2t (73) by restriction cloning (pPL2t.pActA-RFP). To integrate pPL1.pHyper-GFP into Δ*actA* (DPL3078) (74), this strain was cured of its endogenous prophage at the *comK-attBB’* site as previously described (72). Δ*actA* was grown overnight in BHI broth at 37°C with shaking. The next day, 8.5 x 10^8^ U153 phage was combined with 4.25 x 10^7^ bacteria (MOI = 20) in a total of 3 mL BHI containing 5 mM CaCl_2_ and incubated at 37°C with shaking for 75 minutes. The mixture was then diluted 1:100 and 1:10,000 into 2 mL BHI in separate culture tubes. Each tube was incubated at 37°C with shaking for several hours until the 1:100 dilution reached an OD of approximately 0.1. At this point, the 1:10,000 dilution was diluted and spread plated to obtain single colonies. Colonies were screened by PCR with primers specific for *comK-attBB’* for presence of a 417 bp band to indicate successful phage curing. pPL1.pHyper-GFP and pPL2t.pActA-RFP were integrated into phage-cured Δ*actA* via trans-conjugation from *E. coli* SM10, generating the strain designated as GFP-*Lm*.

### Measuring bacterial uptake by macrophages via gentamicin protection assay

Cells were plated in 24-well plates at a density of 5 x 10^5^ (iBMM) or 6 x 10^5^ (BMDM) cells per well the day before infection. 5 x 10^5^ THP-1 cells were plated in the presence of 100 ng/mL phorbol 12-myristate 13-acetate (PMA) 48 hours prior to infection to differentiate cells. Overnight bacterial cultures were diluted in BHI to an OD of 0.05-0.1 and grown at 37°C with shaking to an OD of 1-2. Bacteria were washed twice with PBS, diluted in cell culture medium, and added to cell monolayers at MOI = 1. For experiments comparing different bacterial strains or species, the inoculation medium was serially diluted and plated to enumerate inocula. When indicated, 24-well plates were centrifuged for 3 minutes at 300 x g after adding bacteria to cells. Cells were incubated with bacteria for 30 minutes, washed twice with PBS, and incubated with medium containing gentamicin for an additional 30 minutes. To enumerate intracellular bacteria, cells were washed twice with PBS and lysed in 250 μL cold 0.1% Triton-X in PBS. Lysates were serially diluted and plated for CFU.

### Lentiviral transduction

HEK293T cells were transfected with pVSV-G (Addgene #138479), psPAX2 (Addgene #12260), and a lentiviral vector at a 2:2:1 molar ratio. The following day, culture media was replaced with fresh D-10 supplemented with non-essential amino acids (Cytiva) and 25 mM HEPES. Lentivirus-containing supernatant was collected 48 hours post-transfection, spun at 300 x g for 8 minutes to remove cell debris, aliquoted, and frozen at −80°C. For transductions, cells were plated with lentiviral supernatants in the presence of 8 μg/mL polybrene, and antibiotics were added 48 hours post-transduction. Transduced cells were passaged in the presence of antibiotics for at least 1 week to allow complete selection.

### CRISPR/Cas9 screen

The mouse Brie pooled sgRNA library on the lentiGuide-Puro backbone was obtained from David Root and John Doench (Addgene #73633) (23). The BRIE library contains 78,637 sgRNAs, including 1000 non-targeting controls, and targets 19,674 genes using 4 sgRNAs per gene. The sgRNA library was packaged into lentivirus and transduced into iBMM-Cas9 cells as described above at MOI = 0.3 to ensure single integration events. Transduced cells were selected with puromycin for 1 week and maintained at 1000-fold coverage of the library. Three independent knockout libraries were generated and screened.

Knockout libraries were plated at 1000-fold coverage in 150 mm tissue culture dishes. The GFP-*Lm* strain was grown overnight in BHI broth at 30°C stationary, washed twice with sterile PBS, and used to infect the knockout library at MOI = 10; sterile PBS was added to mock-infected control cells. At 4 hours post-infection cells were washed twice with PBS, and medium containing gentamicin was added to kill extracellular bacteria. At 6 hours post-infection cells were washed twice with PBS, harvested with 0.05% trypsin-EDTA, and fixed with 1% formaldehyde in PBS for 10 min. Fixed cells were washed twice with Flow Buffer (PBS with 5% FBS) and stored at 4°C overnight. The following day, fluorescence-activated cell sorting (FACS) was performed using a BD FACSAria II cell sorter. To isolate uninfected cells, gates were drawn based on mock-infected cells (negative control) and wildtype cells infected with Δ*actA* pPL1.pHyper-GFP pPL2t.pHyper-RFP (positive control). Double negative cells (GFP^-^ RFP^-^) were collected from the infected knockout libraries. Genomic DNA (gDNA) was isolated from mock-infected and sorted populations from three biological replicates as previously described(75). The sgRNA locus was amplified and barcoded by PCR using KAPA HiFi polymerase and Ultramer DNA Oligos from IDT (S3 Table). Enough gDNA was amplified from mock-infected libraries to maintain 1000-fold coverage of the library, using an estimation of 6.6 μg gDNA per 10^6^ cells; for sorted populations, all gDNA was amplified. PCR mixes were split into multiple 50 μL reactions, each containing 5 μg gDNA, and cycled 23 times according to manufacturer recommendations. Amplicons were purified using AMPure XP beads, pooled at 1:1 molar ratios, and sequenced on an Illumina HiSeq2500. sgRNAs enriched in the GFP-negative population were identified using MAGeCK as previously described (24). Gene ontology analysis was performed using Metascape (25).

### Construction of knockout cell lines and CRISPR screen validation by flow cytometry

Individual sgRNAs (2 per gene) from the Brie library or designed using Benchling (S3 Table) were cloned into lentiGuide-Puro (Addgene #52963), lentiCRISPR v2-blast (Addgene #98293), or sgOpti (Addgene #85681) as previously described (22,76–78). Briefly, sgRNAs were annealed, phosphorylated with T4 Polynucleotide Kinase, digested with BsmBI, and ligated into lentiGuide-Puro, lentiCRISPR v2-blast, or sgOpti. Plasmids were packaged into lentivirus and transduced into iBMM-Cas9 cells as described above. To confirm genome-editing, gDNA was isolated using Zymo Quick-DNA Miniprep kits, cut sites were amplified by PCR, and Sanger sequencing performed. Sequences were analyzed using ICE analysis (26) to estimate InDel and knockout efficiency. Clonal cell lines were derived from *Pten* knockout iBMMs (sgRNA1) for further analysis by single-cell sorting.

Knockout cell lines were plated at a density of 10^6^ cells/well in 12-well plates and infected with GFP-*Lm* at MOI = 10. After 4 hours, cells were washed twice with PBS and medium containing gentamicin was added for an additional 2 hours. Cells were then washed twice with PBS, harvested with 0.05% trypsin-EDTA, and fixed with 1% formaldehyde in Flow Buffer for 10 minutes. Samples were washed twice with Flow Buffer and stored in Flow Buffer at 4°C overnight. The next day, samples were analyzed on a BD LSRII flow cytometer. GFP fluorescence was measured and quantified by setting a gate for GFP-positive cells on uninfected cells with 5% background.

### Chemical inhibitors

All chemical inhibitors were added to cells 1 hour prior to infection and kept in the culture medium throughout the experiment. To avoid inhibition of bacterial transcription by actinomycin D, cells pre-treated with actinomycin D were washed twice with PBS immediately prior to infection and medium without inhibitor was used for the remainder of the experiment. Inhibitors had no effect on *Lm* viability, as measured by exposing *Lm* to each inhibitor for 1 hour in D-10 and enumerating CFU. All inhibitors were dissolved in DMSO except bpV(pic) which was dissolved in water. Inhibitors were used at the following concentrations: actinomycin D (Sigma-Aldrich), 10 μg/mL; cycloheximide (Sigma-Aldrich), 1 μg/mL; bpV(pic) (Cayman Chemical Company), 5 μM; CHIR99021 (Cayman Chemical Company), 10 μM; parthenolide (Sigma-Aldrich), 10 μM; wortmannin (Santa Cruz Biotechnology), 100 nM; INK1117 (GlpBio), 1 μM; TGX221 (Cayman Chemical Company), 10 μM; IC87114 (Cayman Chemical Company), 10 μM; and IPI549 (MedChemExpress), 50 nM.

### Phagocytosis of fluorescent beads

Phagocytosis of fluorescence beads was measured using IgG FITC Phagocytosis Assay Kits (Cayman Chemical Company) according to manufacturer instructions. Briefly, FITC-labeled, IgG-opsonized beads were diluted 1:250 in cell culture medium and incubated with iBMMs for 30 minutes. Cells were washed once in assay buffer and incubated in 0.04% Trypan Blue in assay buffer for 2 minutes to quench fluorescence from extracellular beads. Cells were washed once more with assay buffer, harvested using 0.05% trypsin-EDTA, and resuspended in Flow Buffer. Samples were analyzed on a BD LSRII flow cytometer to quantify FITC-positive cells.

### Invasion assays in non-phagocytic cells

Cells were plated at a density of 5 x 10^5^ cells per well in 12-well plates (TIB73) or 2 x 10^5^ cells per well in 24-well plates (Caco-2) the day before infection. Overnight *Lm* cultures were diluted to an OD of 0.05 and grown at 37°C shaking until reaching an OD of 1-2. Bacteria were washed twice with PBS, diluted in cell culture medium, and added to cell monolayers. Cells were infected with MOI = 50 (TIB73) or MOI = 10 (Caco-2). Medium containing 0.1% FBS was used for TIB73 infections and replaced with D-10 for subsequent steps. Cells were incubated with bacteria for 1 hour, washed twice with pre-warmed D-10, and incubated with medium containing gentamicin for an additional 30 minutes. To enumerate intracellular bacteria, cells were washed twice with pre-warmed D-10 and lysed in 250 μL cold 0.1% Triton-X in PBS. Lysates were serially diluted and plated for CFU.

### Immunoblotting

For PTEN immunoblots, 10^6^ iBMMs per well were plated in 12-well plates. The following day, cell lysates were prepared as described below. For pAkt immunoblots, 1.2 x 10^6^ BMDMs per well were plated in 12-well plates. The following day, overnight bacterial cultures were diluted to an OD of 0.05 in BHI and grown at 37°C with shaking to an OD of 1-2. To eliminate background Akt phosphorylation, BMDMs were serum-starved by incubating in FBS- and CSF-free BMDM medium for 2 hours prior to infection. Bacteria were washed twice with PBS, diluted in FBS- and CSF-free BMDM medium, and added to cell monolayers at MOI = 100. At each timepoint, cells were washed twice in PBS and lysed in 50-100 μL cold RIPA buffer containing Halt Protease and Phosphatase Inhibitor Cocktail (Thermo Scientific). Lysates were incubated on ice for 15 minutes and centrifuged at 15,000 x g for 15 minutes at 4°C to pellet cell debris. Protein concentration of clarified lysates was quantified using a Pierce BCA Protein Assay Kit (Thermo Scientific). 30 μg of each sample were separated by SDS-PAGE and transferred to PVD-F membranes. Membranes were blocked for 1 hour at room temperature in Intercept (TBS) Blocking Buffer (LI-COR) for pAkt blots or Intercept (PBS) Blocking Buffer (LI-COR) for PTEN immunoblots. Blocked membranes were incubated overnight at 4°C with anti-Phospho-Akt (Ser473) antibody (Cell Signaling Technology #9271) or anti-PTEN antibody (Cell Signaling Technology #9559) diluted 1:1,000 in TBS antibody buffer (50% TBS, 50% TBS Blocking Buffer) or PBS antibody buffer (50% PBS, 50% PBS Blocking Buffer), respectively. The following day, membranes were washed with TBST and incubated for 1 hour at room temperature with anti-β-Actin antibody (Cell Signaling Technology #3700) diluted 1:1,000 in PBS antibody buffer. Membranes were washed with TBST and incubated with goat anti-rabbit Alexa Fluor 680 (Invitrogen) and goat anti-mouse IRDye 800CW (LI-COR) antibodies diluted 1:5,000 in PBS antibody buffer for 1 hour at room temperature. Blotted membranes were washed a final time with TBST and imaged on a LI-COR Odyssey Fc imager. For pAkt blots, membranes were stripped by incubation at 55°C in stripping buffer (62.5 mM Tris pH 6.8, 2% SDS, 0.8% β-mercaptoethanol) for 45 minutes followed by 5-10 washes in water. Membranes were reblocked with PBS Blocking Buffer and reprobed overnight at 4°C with anti-Akt antibody (Cell Signaling Technology #9272) diluted 1:1,000 in PBS antibody buffer, followed by goat anti-rabbit Alexa Fluor 680 for 1 hour at room temperature.

### Intracellular growth curves

5 x 10^5^ iBMMs per well were plated in 24-well plates overnight. *Lm* cultures were grown overnight at 30°C stationary. The next day, overnight cultures were washed twice with PBS, diluted in cell culture medium, and added to cell monolayers at MOI = 1. After 30 minutes, cells were washed twice and medium containing gentamicin was added to kill extracellular bacteria. At various timepoints (0.5, 2, 5, and 8 hours post-infection), cells were washed twice with PBS and lysed in 250 μL cold 0.1% Triton-X in PBS. Lysates were serially diluted and plated to enumerate intracellular CFU.

### Immunofluorescence microscopy

iBMMs were plated onto 12 mm sterile glass coverslips in 24-well plates at a density of 5 x 10^5^ cells per well. *Lm* constitutively expressing mCherry (pPL2.pHyper-mCherry) (79) was grown overnight in BHI at 37°C shaking. The following day, overnight bacterial cultures were diluted in BHI to an OD of 0.05 and grown at 37°C with shaking to an OD of 1-2. Bacteria were washed twice with PBS, diluted in cell culture medium, and added to cell monolayers at MOI = 5. Plates were centrifuged for 3 minutes at 300 x g to synchronize infection. At each timepoint, coverslips were washed twice with PBS, removed from the plate, and inverted onto 20 μL of 4% formaldehyde on parafilm. Cells were fixed for 10 minutes and washed by dipping into three 50 mL conicals of PBS, 10 times each. Coverslips were washed in this manner in between each of the following incubations. Fixed coverslips were blocked with 1% BSA in PBS (staining buffer) for 10 minutes and stored in staining buffer overnight at 4°C. To block non-specific binding to Fc receptors, coverslips were incubated with mouse Fc Block (BD) diluted 1:50 in staining buffer for 30 minutes. For immunofluorescence staining, coverslips were incubated for 30 minutes with Listeria O Antisera (BD) diluted 1:100 in staining buffer, followed by goat anti-rabbit Alexa Fluor 488 secondary antibody (Invitrogen) diluted 1:200 in staining buffer. Stained coverslips were mounted on glass slides with ProLong Diamond Antifade Mountant with DAPI (Invitrogen) and imaged on a Keyence BZ-X710 fluorescence microscope at 40X magnification.

### Expression of PTEN variants in iBMMs

cDNA was prepared from wildtype iBMMs by extracting total RNA using TRIzol (Invitrogen) and reverse transcription with an iScript cDNA Synthesis Kit (Bio-Rad). PTEN coding sequence was PCR amplified from cDNA and cloned into pLV-EF1a-IRES-Neo (Addgene #85139) (80) using restriction cloning. To prevent CRISPR/Cas9-mediated genome editing in *Pten*^−/−^ iBMMs which constitutively express Cas9 and *Pten*-specific sgRNA, synonymous point mutations were made in each amino acid within the sgRNA recognition site using site-directed mutagenesis to generate a CRISPR-resistant *Pten* allele (PTEN-CR). PTEN-CR was further mutagenized using site-directed mutagenesis to create G129E or G129R point mutations. Empty pLV-EF1a-IRES-Neo, pLV.PTEN-CR, pLV.PTEN-CR-G129E, or pLV.PTEN-CR-G129R were packaged into lentivirus and transduced into *Pten*^−/−^ iBMMs or wildtype iBMMs (empty vector only) as described above. After antibiotic selection, protein expression of each PTEN variant was confirmed by immunoblot as described above.

### Mouse infections

Oral infections were performed as previously described (44,81). 8-10 week old male and female mice were used for infections. 5 mg/mL streptomycin was added to drinking water for 48 hours to eliminate gut flora. Food and water were then removed 16 hours prior to infection to initiate overnight fasting. *Lm* was grown overnight in BHI at 30°C statically. The following day, cultures were diluted 1:10 into 5 mL fresh BHI and grown at 37°C with shaking for 2 hours. Bacteria were washed twice with PBS and 10^8^ CFU in 20 μL was fed to mice via pipette. We found that *Lm* Li2 did not colonize mice as well as 10403S at the same infectious dose, which may be due to the presence of residual streptomycin in the mice at the time of infection (43). To overcome this, we used a higher infectious dose of 5 x 10^8^, which resulted in bacterial burdens similar to 10403S at 5 dpi. Food and water were returned immediately after infection. Bacterial suspensions were serially diluted and plated to enumerate inocula. Mice were humanely euthanized at 1-, 3-, or 5-days post-infection and tissues were collected. Tissues were homogenized in 0.1% NP-40 in the following volumes: MLN, 3 mL; cecum (contents removed and tissue rinsed with PBS), 4 mL; spleen, 3 mL; and liver, 5 mL. Feces were homogenized in 1 mL 0.1% NP-40 with a sterile stick. Gallbladders were ruptured in 500 μL 0.1% NP-40 with a sterile stick. Samples were serially diluted and plated to enumerate CFU in each organ.

### Intestinal tissue fractionation

Fractionation of intestinal tissues was performed as previously described (3,82). Luminal contents of colons and ileums (distal 12 cm of small intestine) were removed by gently squeezing tissues with forceps. Tissues were flushed with 10 mL PBS using a 25 G needle and stored in 15 mL conicals in 5 mL PBS at 4°C overnight. The following day, tissues were cut longitudinally, incubated in 3 mL 6 mM N-acetylcysteine (NAC) for 2 minutes, and shaken vigorously to remove the mucus layer. The NAC wash was repeated twice more for 3 total washes, transferring tissues to fresh tubes each time. To remove epithelial cells, tissues were then cut into small pieces and added to 15 mL conicals with 5 mL RPMI containing 5% FBS, 5 mM EDTA, and 1 mM DTT. Samples were incubated at 37°C with shaking for 20 min, vortexed gently, and filtered through a 70 μm cell strainer. This process was repeated twice more for 3 total EDTA/DTT washes. Tissue pieces were washed once with 10 mL PBS to remove DTT. To isolate lamina propria (LP) cells, tissue pieces were then added to 15 mL conicals with 3 mL RPMI containing 5% FBS, 1 mg/mL collagenase type IV (Sigma-Aldrich), and 40 μg/mL DNase I (Roche). Samples were incubated at 37°C with shaking for 40 min, vortexed, and filtered through 70 μm cell strainers. This process was repeated once more if necessary, until no more large tissue pieces remained. Fractions (NAC washes, EDTA/DTT washes, and LP suspensions) were pooled, centrifuged at 12,000 x g for 20 minutes at 4°C, and pellets resuspended in 150 μL 0.1% NP-40. Samples were serially diluted and plated to enumerate CFU in each fraction.

## References

1. Quereda JJ, Leclercq A, Moura A, Vales G, Gómez-Martín Á, García-Muñoz Á, et al. Listeria valentina sp. nov., isolated from a water trough and the faeces of healthy sheep. International Journal of Systematic and Evolutionary Microbiology. 2020 Nov 1;70(11):5868–79.

2. Vázquez-Boland JA, Kuhn M, Berche P, Chakraborty T, Domínguez-Bernal G, Goebel W, et al. Listeria Pathogenesis and Molecular Virulence Determinants. Clinical Microbiology Reviews. 2001 Jul;14(3):584–640.

3. Bou Ghanem EN, Jones GS, Myers-Morales T, Patil PD, Hidayatullah AN, D’Orazio SEF. InlA Promotes Dissemination of Listeria monocytogenes to the Mesenteric Lymph Nodes during Food Borne Infection of Mice. PLoS Pathog. 2012 Nov 15;8(11):e1003015.

4. Melton-Witt JA, Rafelski SM, Portnoy DA, Bakardjiev AI. Oral Infection with Signature-Tagged Listeria monocytogenes Reveals Organ-Specific Growth and Dissemination Routes in Guinea Pigs. Infection and Immunity. 2011;80(2):720–32.

5. Ireton K, Mortuza R, Gyanwali GC, Gianfelice A, Hussain M. Role of internalin proteins in the pathogenesis of Listeria monocytogenes. Molecular Microbiology. 2021;116(6):1407–19.

6. Jones GS, Bussell KM, Myers-Morales T, Fieldhouse AM, Bou Ghanem EN, D’Orazio SEF. Intracellular Listeria monocytogenes Comprises a Minimal but Vital Fraction of the Intestinal Burden following Foodborne Infection. Infect Immun. 2015 Aug;83(8):3146–56.

7. Drevets DA. Dissemination of Listeria monocytogenesby Infected Phagocytes. Infection and Immunity. 1999;67(7):3512–7.

8. Drevets DA, Jelinek TA, Freitag NE. Listeria monocytogenes-Infected Phagocytes Can Initiate Central Nervous System Infection in Mice. Infection and Immunity. 2001;69(3):1344–50.

9. Drevets DA, Dillon MJ, Schawang JS, van Rooijen N, Ehrchen J, Sunderkötter C, et al. The Ly-6C ^high^ Monocyte Subpopulation Transports *Listeria monocytogenes* into the Brain during Systemic Infection of Mice. J Immunol. 2004 Apr 1;172(7):4418–24.

10. Join-Lambert OF, Ezine S, Le Monnier A, Jaubert F, Okabe M, Berche P, et al. Listeria monocytogenes-infected bone marrow myeloid cells promote bacterial invasion of the central nervous system. Cellular Microbiology. 2005;7(2):167–80.

11. Maudet C, Kheloufi M, Levallois S, Gaillard J, Huang L, Gaultier C, et al. Bacterial inhibition of Fas-mediated killing promotes neuroinvasion and persistence. Nature. 2022 Mar;603(7903):900–6.

12. Pamer EG. Immune responses to Listeria monocytogenes. Nat Rev Immunol. 2004 Oct;4(10):812–23.

13. Unanue ER. Inter-relationship among macrophages, natural killer cells and neutrophils in early stages of Listeria resistance. Current Opinion in Immunology. 1997 Feb 1;9(1):35–43.

14. Buchmeier NA, Schreiber RD. Requirement of endogenous interferon-gamma production for resolution of Listeria monocytogenes infection. Proceedings of the National Academy of Sciences. 1985 Nov;82(21):7404–8.

15. Dai WJ, Bartens W, Köhler G, Hufnagel M, Kopf M, Brombacher F. Impaired macrophage listericidal and cytokine activities are responsible for the rapid death of Listeria monocytogenes-infected IFN-gamma receptor-deficient mice. The Journal of Immunology. 1997 Jun 1;158(11):5297–304.

16. Portnoy DA, Schreiber RD, Connelly P, Tilney LG. Gamma interferon limits access of Listeria monocytogenes to the macrophage cytoplasm. Journal of Experimental Medicine. 1989 Dec 1;170(6):2141–6.

17. Portnoy DA, Auerbuch V, Glomski IJ. The cell biology of Listeria monocytogenes infection. J Cell Biol. 2002 Aug 5;158(3):409–14.

18. Drevets DA, Leenen PJM, Campbell PA. Complement Receptor Type 3 Mediates Phagocytosis and Killing ofListeria monocytogenesby a TNF-α- and IFN-γ-Stimulated Macrophage Precursor Hybrid. Cellular Immunology. 1996 Apr 10;169(1):1–6.

19. Sorbara MT, Foerster EG, Tsalikis J, Abdel-Nour M, Mangiapane J, Sirluck-Schroeder I, et al. Complement C3 Drives Autophagy-Dependent Restriction of Cyto-invasive Bacteria. Cell Host & Microbe. 2018 May 9;23(5):644–652.e5.

20. Drevets DA, Canono BP, Campbell PA. Listericidal and nonlistericidal mouse macrophages differ in complement receptor type 3-mediated phagocytosis of L. monocytogenes and in preventing escape of the bacteria into the cytoplasm. J Leukoc Biol. 1992 Jul;52(1):70–9.

21. Pierce MM, Gibson RE, Rodgers FG. Opsonin-independent adherence and phagocytosis of Listeria monocytogenes by murine peritoneal macrophages. Journal of Medical Microbiology. 1996 Oct 1;45(4):258–62.

22. Kiritsy MC, Ankley LM, Trombley J, Huizinga GP, Lord AE, Orning P, et al. A genetic screen in macrophages identifies new regulators of IFNγ-inducible MHCII that contribute to T cell activation. Horng T, Garrett WS, Divangahi M, Zanoni I, editors. eLife. 2021 Nov 8;10:e65110.

23. Doench JG, Fusi N, Sullender M, Hegde M, Vaimberg EW, Donovan KF, et al. Optimized sgRNA design to maximize activity and minimize off-target effects of CRISPR-Cas9. Nat Biotechnol. 2016 Feb;34(2):184–91.

24. Li W, Xu H, Xiao T, Cong L, Love MI, Zhang F, et al. MAGeCK enables robust identification of essential genes from genome-scale CRISPR/Cas9 knockout screens. Genome Biology. 2014 Dec 5;15(12):554.

25. Zhou Y, Zhou B, Pache L, Chang M, Khodabakhshi AH, Tanaseichuk O, et al. Metascape provides a biologist-oriented resource for the analysis of systems-level datasets. Nat Commun. 2019 Apr 3;10:1523.

26. Hsiau T, Conant D, Rossi N, Maures T, Waite K, Yang J, et al. Inference of CRISPR Edits from Sanger Trace Data. bioRxiv. 2019 Aug 10;251082.

27. Keniry M, Parsons R. The role of PTEN signaling perturbations in cancer and in targeted therapy. Oncogene. 2008 Sep;27(41):5477–85.

28. Maehama T, Dixon JE. The Tumor Suppressor, PTEN/MMAC1, Dephosphorylates the Lipid Second Messenger, Phosphatidylinositol 3,4,5-Trisphosphate *. Journal of Biological Chemistry. 1998 May 29;273(22):13375–8.

29. Stambolic V, Suzuki A, de la Pompa JL, Brothers GM, Mirtsos C, Sasaki T, et al. Negative Regulation of PKB/Akt-Dependent Cell Survival by the Tumor Suppressor PTEN. Cell. 1998 Oct 2;95(1):29–39.

30. Sharma L, Wu W, Dholakiya SL, Gorasiya S, Wu J, Sitapara R, et al. Assessment of Phagocytic Activity of Cultured Macrophages Using Fluorescence Microscopy and Flow Cytometry. In: Vancurova I, editor. Cytokine Bioassays: Methods and Protocols [Internet]. New York, NY: Springer; 2014 [cited 2022 Sep 28]. p. 137–45. (Methods in Molecular Biology). Available from: https://doi.org/10.1007/978-1-4939-0928-5_12

31. Schmid AC, Byrne RD, Vilar R, Woscholski R. Bisperoxovanadium compounds are potent PTEN inhibitors. FEBS Lett. 2004 May 21;566(1–3):35–8.

32. Ireton K. Entry of the bacterial pathogen Listeria monocytogenes into mammalian cells. Cellular Microbiology. 2007;9(6):1365–75.

33. Myers MP, Pass I, Batty IH, Van der Kaay J, Stolarov JP, Hemmings BA, et al. The lipid phosphatase activity of PTEN is critical for its tumor supressor function. Proceedings of the National Academy of Sciences. 1998 Nov 10;95(23):13513–8.

34. Currie RA, Walker KS, Gray A, Deak M, Casamayor A, Downes CP, et al. Role of phosphatidylinositol 3,4,5-trisphosphate in regulating the activity and localization of 3-phosphoinositide-dependent protein kinase-1. Biochemical Journal. 1999 Jan 25;337(3):575–83.

35. Bilanges B, Posor Y, Vanhaesebroeck B. PI3K isoforms in cell signalling and vesicle trafficking. Nat Rev Mol Cell Biol. 2019 Sep;20(9):515–34.

36. Arbibe L, Mira JP, Teusch N, Kline L, Guha M, Mackman N, et al. Toll-like receptor 2–mediated NF-κB activation requires a Rac1-dependent pathway. Nat Immunol. 2000 Dec;1(6):533–40.

37. Rhee SH, Kim H, Moyer MP, Pothoulakis C. Role of MyD88 in Phosphatidylinositol 3-Kinase Activation by Flagellin/Toll-like Receptor 5 Engagement in Colonic Epithelial Cells *. Journal of Biological Chemistry. 2006 Jul 7;281(27):18560–8.

38. Hayashi F, Smith KD, Ozinsky A, Hawn TR, Yi EC, Goodlett DR, et al. The innate immune response to bacterial flagellin is mediated by Toll-like receptor 5. Nature. 2001 Apr;410(6832):1099–103.

39. Torres D, Barrier M, Bihl F, Quesniaux VJF, Maillet I, Akira S, et al. Toll-Like Receptor 2 Is Required for Optimal Control of Listeria monocytogenes Infection. Infect Immun. 2004 Apr;72(4):2131–9.

40. Cesinger MR, Daramola OI, Kwiatkowski LM, Reniere ML. The Transcriptional Regulator SpxA1 Influences the Morphology and Virulence of Listeria monocytogenes. Infection and Immunity. 2022 Sep 14;90(10):e00211–22.

41. Gründling A, Burrack LS, Bouwer HGA, Higgins DE. Listeria monocytogenes regulates flagellar motility gene expression through MogR, a transcriptional repressor required for virulence. Proceedings of the National Academy of Sciences. 2004 Aug 17;101(33):12318–23.

42. Kim SH, Park MK, Kim JY, Chuong PD, Lee YS, Yoon BS, et al. Development of a sandwich ELISA for the detection of Listeria spp. using specific flagella antibodies. J Vet Sci. 2019 Feb 11;6(1):41–6.

43. Becattini S, Littmann ER, Carter RA, Kim SG, Morjaria SM, Ling L, et al. Commensal microbes provide first line defense against Listeria monocytogenes infection. Journal of Experimental Medicine. 2017 Jun 6;214(7):1973–89.

44. Louie A, Zhang T, Becattini S, Waldor MK, Portnoy DA. A Multiorgan Trafficking Circuit Provides Purifying Selection of Listeria monocytogenes Virulence Genes. Miller SI, editor. mBio. 2019 Dec 24;10(6):e02948–19.

45. Gillooly DJ, Simonsen A, Stenmark H. Phosphoinositides and phagocytosis. J Cell Biol. 2001 Oct 1;155(1):15–8.

46. Lv Y, Fang L, Ding P, Liu R. PI3K/Akt-Beclin1 signaling pathway positively regulates phagocytosis and negatively mediates NF-κB-dependent inflammation in Staphylococcus aureus-infected macrophages. Biochemical and Biophysical Research Communications. 2019 Mar 5;510(2):284–9.

47. Tachado SD, Samrakandi MM, Cirillo JD. Non-Opsonic Phagocytosis of Legionella pneumophila by Macrophages Is Mediated by Phosphatidylinositol 3-Kinase. PLOS ONE. 2008 Oct 2;3(10):e3324.

48. Lovewell RR, Hayes SM, O’Toole GA, Berwin B. Pseudomonas aeruginosa flagellar motility activates the phagocyte PI3K/Akt pathway to induce phagocytic engulfment. American Journal of Physiology-Lung Cellular and Molecular Physiology. 2014 Apr;306(7):L698–707.

49. Shin OS, Miller LS, Modlin RL, Akira S, Uematsu S, Hu LT. Downstream Signals for MyD88-Mediated Phagocytosis of *Borrelia burgdorferi* Can Be Initiated by TRIF and Are Dependent on PI3K. J Immunol. 2009 Jul 1;183(1):491–8.

50. Hubbard LLN, Wilke CA, White ES, Moore BB. PTEN Limits Alveolar Macrophage Function against Pseudomonas aeruginosa after Bone Marrow Transplantation. Am J Respir Cell Mol Biol. 2011 Nov 1;45(5):1050–8.

51. Schabbauer G, Matt U, Günzl P, Warszawska J, Furtner T, Hainzl E, et al. Myeloid PTEN Promotes Inflammation but Impairs Bactericidal Activities during Murine Pneumococcal Pneumonia. J Immunol. 2010 Jul 1;185(1):468–76.

52. Serezani CH, Kane S, Medeiros AI, Cornett AM, Kim SH, Marques MM, et al. PTEN Directly Activates the Actin Depolymerization Factor Cofilin-1 During PGE2-Mediated Inhibition of Phagocytosis of Fungi. Sci Signal. 2012 Feb 7;5(210):ra12–ra12.

53. Mondal S, Ghosh-Roy S, Loison F, Li Y, Jia Y, Harris C, et al. PTEN Negatively Regulates Engulfment of Apoptotic Cells by Modulating Activation of Rac GTPase. The Journal of Immunology. 2011 Dec 1;187(11):5783–94.

54. Ireton K, Payrastre B, Chap H, Ogawa W, Sakaue H, Kasuga M, et al. A Role for Phosphoinositide 3-Kinase in Bacterial Invasion. Science. 1996 Nov 1;274(5288):780–2.

55. Czech MP. PIP2 and PIP3: Complex Roles at the Cell Surface. Cell. 2000 Mar 17;100(6):603–6.

56. Mandal K. Review of PIP2 in Cellular Signaling, Functions and Diseases. Int J Mol Sci. 2020 Nov 6;21(21):8342.

57. Perelman SS, Abrams ME, Eitson JL, Chen D, Jimenez A, Mettlen M, et al. Cell-Based Screen Identifies Human Interferon-Stimulated Regulators of Listeria monocytogenes Infection. PLOS Pathogens. 2016 Dec 21;12(12):e1006102.

58. Ishiguro T, Naito M, Yamamoto T, Hasegawa G, Gejyo F, Mitsuyama M, et al. Role of Macrophage Scavenger Receptors in Response to Listeria monocytogenes Infection in Mice. The American Journal of Pathology. 2001 Jan 1;158(1):179–88.

59. Sumrall ET, Keller AP, Shen Y, Loessner MJ. Structure and function of Listeria teichoic acids and their implications. Molecular Microbiology. 2020;113(3):627–37.

60. Sumrall ET, Shen Y, Keller AP, Rismondo J, Pavlou M, Eugster MR, et al. Phage resistance at the cost of virulence: Listeria monocytogenes serovar 4b requires galactosylated teichoic acids for InlB-mediated invasion. PLOS Pathogens. 2019 Oct 7;15(10):e1008032.

61. Disson O, Blériot C, Jacob JM, Serafini N, Dulauroy S, Jouvion G, et al. Peyer’s patch myeloid cells infection by Listeria signals through gp38+ stromal cells and locks intestinal villus invasion. Journal of Experimental Medicine. 2018 Oct 24;215(11):2936–54.

62. McElroy DS, Ashley TJ, D’Orazio SEF. Lymphocytes serve as a reservoir for Listeria monocytogenes growth during infection of mice. Microb Pathog. 2009 Apr;46(4):214–21.

63. Jones GS, D’Orazio SEF. Monocytes Are the Predominant Cell Type Associated with *Listeria monocytogenes* in the Gut, but They Do Not Serve as an Intracellular Growth Niche. J Immunol. 2017 Apr 1;198(7):2796–804.

64. Jones GS, Smith VC, D’Orazio SEF. *Listeria monocytogenes* Replicate in Bone Marrow–Derived CD11c ^+^ Cells but Not in Dendritic Cells Isolated from the Murine Gastrointestinal Tract. J Immunol. 2017 Dec 1;199(11):3789–97.

65. D’Orazio SEF. Innate and Adaptive Immune Responses during Listeria monocytogenes Infection. Microbiology Spectrum [Internet]. 2019 [cited 2021 Jun 18];7(3). Available from: https://journals.asm.org/doi/abs/10.1128/microbiolspec.GPP3-0065-2019

66. Pitts MG, D’Orazio SEF. A Comparison of Oral and Intravenous Mouse Models of Listeriosis. Pathogens. 2018 Mar;7(1):13.

67. Uribe-Querol E, Rosales C. Phagocytosis: Our Current Understanding of a Universal Biological Process. Front Immunol. 2020 Jun 2;11:1066.

68. Lim JJ, Grinstein S, Roth Z. Diversity and Versatility of Phagocytosis: Roles in Innate Immunity, Tissue Remodeling, and Homeostasis. Frontiers in Cellular and Infection Microbiology [Internet]. 2017 [cited 2022 Nov 2];7. Available from: https://www.frontiersin.org/articles/10.3389/fcimb.2017.00191

69. McFarland AP, Burke TP, Carletti AA, Glover RC, Tabakh H, Welch MD, et al. RECON-Dependent Inflammation in Hepatocytes Enhances Listeria monocytogenes Cell-to-Cell Spread. mBio. 2018 May 15;9(3):e00526–18.

70. Young L, Sung J, Stacey G, Masters JR. Detection of Mycoplasma in cell cultures. Nat Protoc. 2010 May;5(5):929–34.

71. Portnoy DA, Jacks PS, Hinrichs DJ. Role of hemolysin for the intracellular growth of Listeria monocytogenes. J Exp Med. 1988 Apr 1;167(4):1459–71.

72. Lauer P, Chow MYN, Loessner MJ, Portnoy DA, Calendar R. Construction, Characterization, and Use of Two Listeria monocytogenes Site-Specific Phage Integration Vectors. Journal of Bacteriology. 2002 Aug;184(15):4177–86.

73. Whiteley AT, Pollock AJ, Portnoy DA. The PAMP c-di-AMP Is Essential for Listeria monocytogenes Growth in Rich but Not Minimal Media due to a Toxic Increase in (p)ppGpp. Cell Host & Microbe. 2015 Jun;17(6):788–98.

74. Skoble J, Portnoy DA, Welch MD. Three Regions within Acta Promote Arp2/3 Complex-Mediated Actin Nucleation and Listeria monocytogenes Motility. Journal of Cell Biology. 2000 Aug 7;150(3):527–38.

75. Chen S, Sanjana NE, Zheng K, Shalem O, Lee K, Shi X, et al. Genome-wide CRISPR Screen in a Mouse Model of Tumor Growth and Metastasis. Cell. 2015 Mar 12;160(6):1246–60.

76. Fulco CP, Munschauer M, Anyoha R, Munson G, Grossman SR, Perez EM, et al. Systematic mapping of functional enhancer-promoter connections with CRISPR interference. Science. 2016 Nov 11;354(6313):769–73.

77. Sanjana NE, Shalem O, Zhang F. Improved vectors and genome-wide libraries for CRISPR screening. Nat Methods. 2014 Aug;11(8):783–4.

78. Stringer BW, Day BW, D’Souza RCJ, Jamieson PR, Ensbey KS, Bruce ZC, et al. A reference collection of patient-derived cell line and xenograft models of proneural, classical and mesenchymal glioblastoma. Sci Rep. 2019 Mar 20;9:4902.

79. Vincent WJB, Freisinger CM, Lam P ying, Huttenlocher A, Sauer JD. Macrophages mediate flagellin induced inflammasome activation and host defense in zebrafish. Cellular Microbiology. 2016;18(4):591–604.

80. Hayer A, Shao L, Chung M, Joubert LM, Yang HW, Tsai FC, et al. Engulfed cadherin fingers are polarized junctional structures between collectively migrating endothelial cells. Nat Cell Biol. 2016 Dec;18(12):1311–23.

81. Halsey CR, Glover RC, Thomason MK, Reniere ML. The redox-responsive transcriptional regulator Rex represses fermentative metabolism and is required for Listeria monocytogenes pathogenesis. PLoS Pathog. 2021 Aug 16;17(8):e1009379.

82. Bou Ghanem EN, Myers-Morales T, D’Orazio SEF. A mouse model of food borne Listeria monocytogenes infection. Curr Protoc Microbiol. 2013 Nov 5;31:9B.3.1-9B.3.16.

